# Flexible and cost-effective genomic surveillance of *P. falciparum* malaria with targeted nanopore sequencing

**DOI:** 10.1101/2023.02.06.527333

**Authors:** Mariateresa de Cesare, Mulenga Mwenda, Anna E. Jeffreys, Jacob Chirwa, Chris Drakeley, Kammerle Schneider, Isaac Ghinai, George B. Busby, Busiku Hamainza, Moonga Hawela, Daniel J. Bridges, Jason A. Hendry

## Abstract

Genomic surveillance of *Plasmodium falciparum* malaria can provide policy-relevant information about antimalarial drug resistance, rapid diagnostic test failure, and the evolution of vaccine targets. Yet the large and low complexity genome of *P. falciparum* limits the scope of genomic surveillance, as whole-genome sequencing approaches are costly and targeted approaches are challenging to develop. Moreover, the majority of the morbidity and mortality caused by *P. falciparum* occurs in sub-Saharan Africa, where resource constraints can make implementing genomic surveillance difficult. Here, we demonstrate a flexible and cost-effective approach for targeted nanopore sequencing of *P. falciparum* to enable genomic surveillance of malaria in low-resource settings. We release open-source software that facilitates rapid and flexible design of amplicon sequencing panels for *P. falciparum*, coupled with a simple and cost-effective protocol that uses dried blood spots as input. We use this software to design two amplicon panels. The first, called NOMADS8, targets seven major antimalarial drug-resistance associated genes as well as the highly polymorphic gene *msp2*. The second, NOMAD16, incorporates an additional eight genes including the vaccine target *csp* and genes coding for the antigens detected in rapid diagnostic tests, *hrp2* and *hrp3*. The panels generate reads between 3 to 4kbp that span the entire coding sequence of most target genes. We validate the panels and protocol on mock and field samples, demonstrating robust sequencing coverage across targets, high single-nucleotide polymorphism calling accuracy within coding sequences, and the ability to explore the within-sample diversity of mixed *P. falciparum* infections.

## Introduction

The malaria parasite species *Plasmodium falciparum* is an example of both the potential value of genomic surveillance and the obstacles that can impede its implementation. Although a variety of antimalarial drugs exist, the evolution of resistance has compromised their efficacy^1, 2^. Most critical is resistance to artemisinin, the dominant chemotherapeutic agent in artemisinin-based combination therapy (ACT) and the foundation of global guidelines for the treatment of malaria^3^. Formerly confined to the Greater Mekong Subregion^4–6^, genetic mutations associated with artemisinin resistance have recently been detected in Uganda^7^ and Rwanda^8^, escalating the risk of ACT failure in sub-Saharan Africa. Additionally, *P. falciparum* parasites with deletions causing false negative rapid diagnostic test (RDT) results have been detected at high frequency in Ghana^9^, Eritrea^10, 11^ and Ethiopia^12, 13^. The causal mutations underlying these phenotypes^13–17^, as well as other common antimalarial drug-resistance phenotypes, are well characterised. By informing on the frequency and distribution of these mutations, genomic surveillance could play a crucial role crafting evidence-based policies to limit their spread and improve malaria control.

Despite its potential value, multiple challenges limit widespread genomic surveillance of *P. falciparum* malaria. First, the nuclear genome is 23Mbp^18^ – considerably larger than typical bacterial (~3-5Mbp)^19, 20^ or viral genomes (~10-100kbp)^21^. At present, this renders whole-genome sequencing (WGS) strategies prohibitively costly to scale. Second, although targeted sequencing strategies — such as those employing multiplex polymerase chain reaction (PCR)^22–24^, molecular-inversion probes (MIP)^25, 26^ or hybrid capture — can be potentially more cost-effective, the genome of *P. falciparum* is extremely (A+T)-rich^18^ and often there is little unique and biochemically-suitable sequence (e.g. for primer or probe design) within proximity of targets. This makes the development of these approaches uniquely difficult for *P. falciparum*. Third, many regions with a high unmet need for *P. falciparum* genomic surveillance are in sub-Saharan Africa, yet most existing targeted sequencing approaches have been developed for Illumina platforms^22–24, 26^. Due to their complexity, costs and maintenance requirements, these platforms are concentrated in centralised sequencing facilities – few of which are in sub-Saharan Africa. Although this situation is improving^27^, deficits in local sequencing capacity still impel many small- and medium-sized labs to ship samples internationally for sequencing. This reduces country engagement, introduces ethical and logistical issues around sample export, and inevitably increases time to result, potentially delaying evidence-based policy decisions.

At the same time, there has been growing use of nanopore sequencing for pathogen genomic surveillance, facilitated by the small and portable MinION sequencing device (Oxford Nanopore Technologies). The MinION can be deployed in low resource settings, requires no maintenance, and permits real-time data analysis^28^. It has been successfully deployed during Ebola^29^ Zika^30^, and SARS-CoV-2 outbreaks^27^. A key advantage of nanopore-based sequencing is the generation of long reads (kbps to Mbps)^31^ which can improve mapping and structural variant detection^32^, while a disadvantage is a higher base-level error rate compared to instruments from Illumina or Pacific Biosciences (PacBio). Although important proof-of-principle studies have demonstrated the feasibility of nanopore-based sequencing of *P. falciparum*, and investigated the consequences of its higher error rate^33, 34^, comparatively little effort has been made to develop methods for routine nanopore-based genomic surveillance of malaria.

In this study, we developed a flexible and cost-effective approach to targeted *P. falciparum* sequencing using the MinION. Flexibility is created through the development of open-source software, called `multiply`, that enables multiplex PCR design for a user-defined set of target genes and/or regions across the *P. falciparum* genome. We use this software to create eight- and sixteen-target amplicon sequencing panels, which encompass genes associated with antimalarial drug resistance, RDT failure, complexity of infection inference and malaria vaccine target *csp*^35, 36^. To sequence these panels we devised an optimised protocol that utilises DBS as input and costs approximately USD $25 per sample. We validated this approach on a set of mock and Zambian field samples collected as DBS, and demonstrate adequate coverage of target genes and a high SNP calling accuracy within coding sequence (CDS). Finally, we show how signals of *P. falciparum* within-sample diversity are present within long-read data through analysis of the surface antigen gene *msp2*.

## Results

### Designing amplicon panels for *P. falciparum* with multiply

New amplicon sequencing panels require the development of a multiplex PCR which, even for a moderate number of targets, entails evaluating vast combinations of primers for off-target binding, primer dimers, or polymorphic sites in the study population. To facilitate this process for amplicon panels where the targets are distributed across larger genomes (i.e. in contrast to tiling PCR of smaller pathogen genomes^37^), we developed software called `multiply` (Fig. 1a). `multiply` provides a rapid and flexible approach to multiplex PCR design given a user-supplied list of target genes and/or regions. Briefly, `multiply` first generates a diverse set of candidate primers for each target using `primer3`^38^. It then searches for polymorphic sites within primer binding locations by intersecting them with user-supplied Variant Call Format (VCF) files; computes primer-dimer scores for all candidate primer pairs using an algorithm similar to that described by Johnston *et al*.^39^; and identifies potential off-target binding sites using `blastn` against the reference genome^40, 41^. Results from these three steps are combined into a cost-function that scores multiplex PCR primer combinations, with a lower score indicating a better predicted performance. Finally, the cost-function is minimised using a greedy search algorithm to identify optimal combinations of primers for the specified targets.

**Figure 1.**
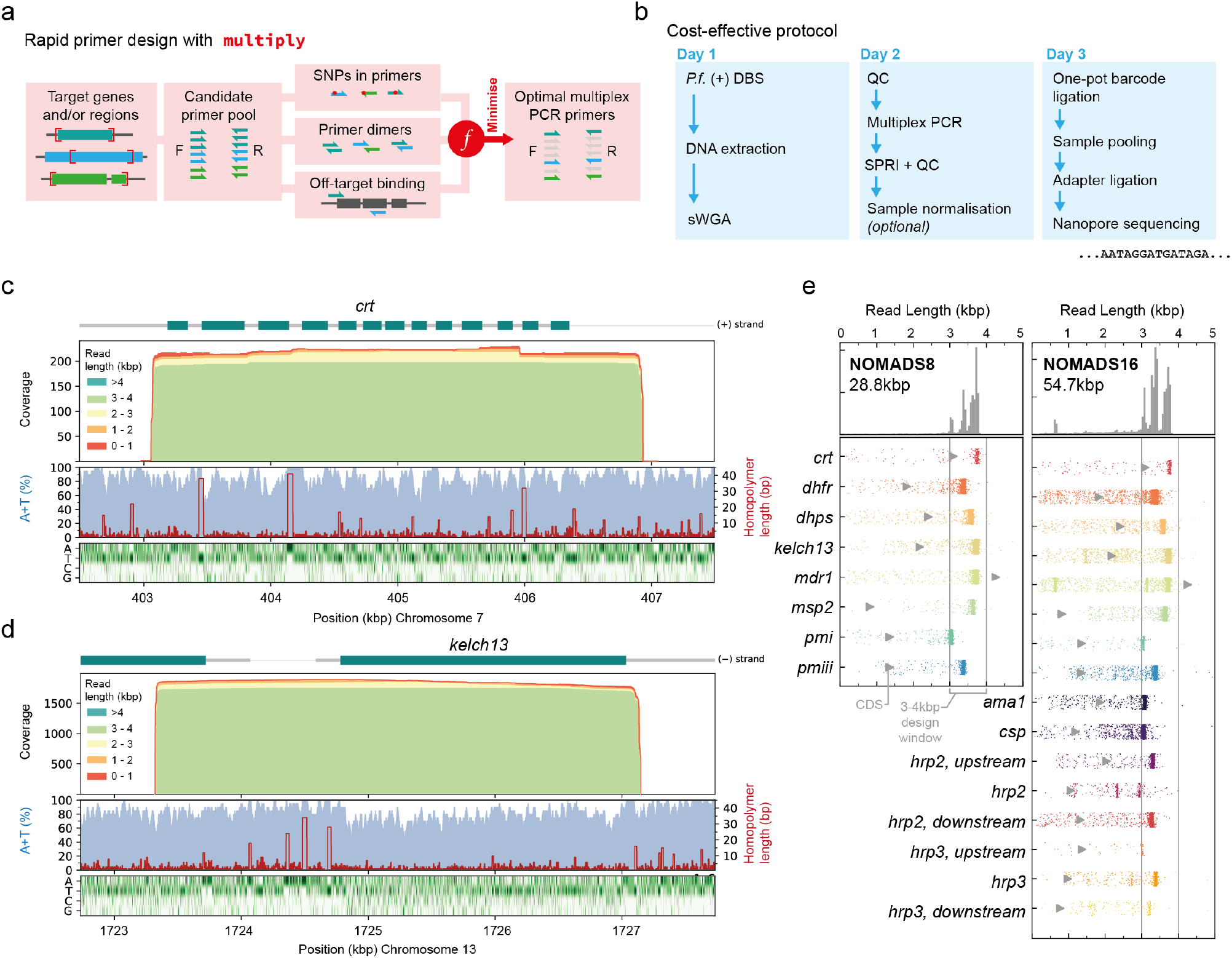
Software-aided design of long-range multiplex PCRs for the low-complexity *P. falciparum* genome. (a) Multiplex PCR primer design workflow by `multiply`. An optimal set of primers is selected from a large candidate pool; minimising SNPs in primer binding sites, primer dimers, and off-target primer binding with a cost function. (b) Schematic of a cost-effective protocol for targeted nanopore sequencing of *P. falciparum* malaria from dried blood spots (DBS) that takes three days and costs ~USD $25 per sample. (c) Histograms of *crt* and (d) *kelch13* coverage stratified by read length. (A+T) percentage in 20bp sliding widows (blue) and homopolymer run length (red) are shown, as well as a heatmap of nucleotide composition. For both genes the entire coding sequence (CDS) is covered in the majority of reads. (e) Read length distributions for NOMADS8 (left, 28.8 kbp total) and NOMADS16 (right, 54.7 kbp total) amplicon panels. Grey triangle indicates coding sequence (CDS) length. Amplicons were designed to be 3-4kbp. Marginal distribution for all amplicons displayed at top. Data for panels (c), (d) and (e) are from a mock sample created from *P. falciparum* 3D7 and human DNA (Methods).

We used `multiply` to develop a multiplex PCR for *P. falciparum* malaria, selecting eight target genes that would maximise the public health utility of our data (Table 1). To leverage the long-read capability of nanopore sequencing, we restricted candidate amplicons to 3 - 4kbp; aiming to produce CDS-spanning amplicons that would still being feasible for PCR. In the design process, `multiply` considered a total of 194 candidate primers across the eight targets. For these candidate primers, it identified 383 high scoring off-target complementary matches in the 3D7 reference genome (> 12bp aligned from the 3’ end). Overall, 209 matches involved candidate forward primers for *dhps*; a candidate reverse primer for *plasmepsin I* (*pmI*) had 35 matches; a candidate reverse primer for *kelch13* had 24 matches; and most other candidate primers had 5 or less matches. By comparing to the 7,113 *P. falciparum* whole genome sequences in the Pf6 data release, `multiply` identified 11 common SNPs (minor allele frequency > 5% in any Pf6 population) within candidate primer binding locations. Of the 18,915 unique pairwise alignments `multiply` computed between candidate primers, 585 had potentially problematic dimer scores (score < −6). Using a greedy search algorithm, `multiply` heuristically minimised these factors to suggest a multiplex PCR primer combination from the over 370 million possibilities given the candidate primer set.

We call the amplicons produced from this multiplex PCR the NOMADS8 (**N**MEC-**O**xford **M**alaria **A**mplicon **D**rug-resistance **S**equencing) panel. In total, the amplicons span 28.8kbp with an (A+T)-composition of 79%. The full coding sequences for 7 of 8 gene targets are included completely within their amplicons. *mdr1* has a coding sequence covering 4259bp; our amplicon is only 3773bp but includes important drug-resistance mutations (e.g. N86Y to D1246Y)^42^. Using PCR conditions with reduced annealing and extension temperatures^43^, we were able to obtain robust amplification of all individual targets and produce bands consistent with expectation for the multiplex, as assessed by agarose gel electrophoresis (Supplementary Fig. 1).

We next used `multiply` to expand the NOMADS8 panel to include an additional eight targets. These included *ama1*, a highly polymorphic gene used in complexity of infection (COI) estimation^44^; the RTS,S and R21 vaccine target *csp*^35, 36^; and the RDT antigen genes *hrp2* and *hrp3*^17^, as well as their flanking genes. To incorporate these eight targets, `multiply` considered an additional 214 candidate primers and, keeping the 16 primers of the NOMADS8 panel fixed, repeated the selection process described above. The resulting amplicons span a total of 54.7kbp and are called the NOMADS16 panel (Table 1).

**Table 1.**
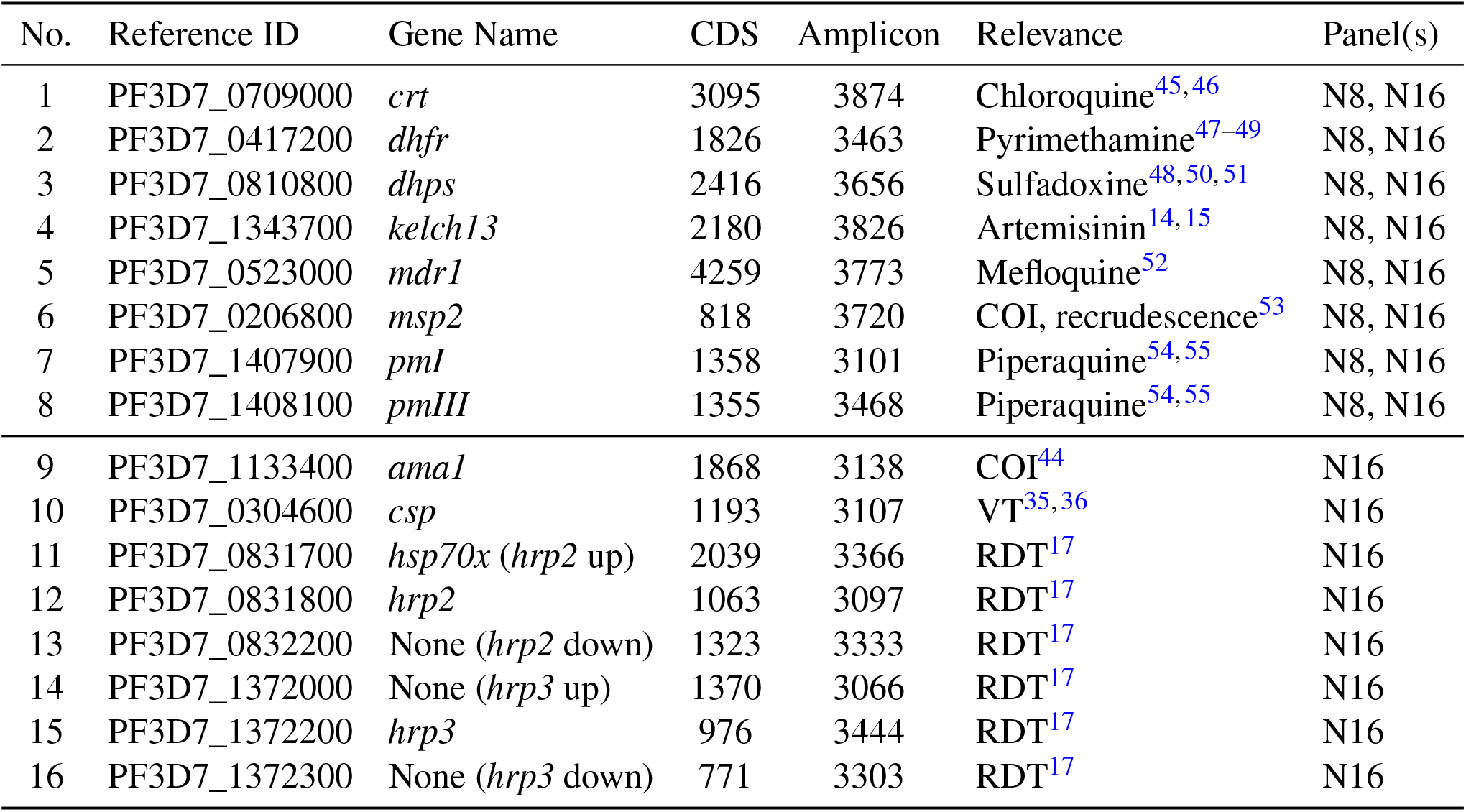
Target genes for the NOMADS8 and NOMADS16 amplicon sequencing panels. For each target, columns give information about the coding-sequence (CDS) and amplicon lengths in base pairs (bp), the epidemiological relevance (COI, complexity of infection; VT, vaccine target) and whether the target is in both the NOMADS8 (N8) and/or the NOMADS16 (N16) panel. The CDS length is measured as the distance between the start and stop codon, including intronic regions if present. ‘*hrp2* up’ and ‘*hrp3* up’ refer to targets upstream of *hrp2* and *hrp3*, respectively; wheres ‘*hrp2* down’ and ‘*hrp3* down’ refer to downstream targets.

### Minimising *P. falciparum* amplicon sequencing costs on the MinION

We combined existing and novel optimisations to minimise the costs of *P. falciparum* amplicon sequencing on the MinION (Fig. 1b). Briefly, our protocol starts with DBS as input for DNA extraction, which are relatively non-invasive and easy to collect. Bulk *P. falciparum* DNA is enriched with a reduced-volume selective-whole genome amplification (sWGA) step, saving approximately USD $4 per sample while still maintaining sufficient yield for subsequent multiplex PCR (Supplementary Fig. 2). Amplicons are barcoded and pooled using a modified version of a simple and cost-effective one-pot protocol^56^. Overall, the protocol can be completed in 2-3 days at USD $25 per sample, assuming 96 samples are run on a R9.4.1 MinION Flow Cell (FLO-MIN106D) without washing (Supplementary Table 1). Smaller batches of 24 samples run on a Flongle Flow Cell (FLO-FLG001) cost a negligible USD $1 extra per sample.

### Producing long-read data for policy-relevant *P. falciparum* genes

We explored the read lengths that are generated with our amplicon panels and protocol by sequencing a mock sample, created by combining *P. falciparum* 3D7 and human DNA *in vitro*, on a Flongle Flow Cell (Methods, Fig. 1c,d,e). For the NOMADS8 panel, the median length of reads that mapped to the *P. falciparum* reference genome and overlapped a target gene was 3.59kbp. All eight target genes had a median read length greater than 3.04kbp and, excluding *mdr1*, on average 91.7% of reads that overlapped a target gene spanned its entire CDS. This included reads spanning all 13 exons of *crt1* and the entire CDS of the artemisinin-resistance associated gene *kelch13* (Fig. 1c,d). In several cases, longer amplicons enabled `multiply` to select primers that bind to regions with more moderate (A+T) compositions in adjacent genes, and this was the case for the forward primer used to amplify *kelch13* (Fig 1d). Similarly, the median length of target-overlapping reads for the NOMADS16 panel was 3.37kbp, with an average of 88.2% of these completely spanning their target’s CDS (excluding *mdr1*).

### Characterising sequencing efficiency and coverage across mock and field samples

Sufficient coverage over target regions is a precondition for accurate variant calling and other downstream analyses. Whether this is achieved depends on the total sequencing throughput, the proportion of that throughput that is on-target (i.e. maps to the intended organism and regions), and how uniformly on-target throughput is distributed across the target regions and samples.

We characterised the coverage generated by our protocol by running experiments with both the NOMADS8 and NOMADS16 panels on two different sample sets. The first set included 24 mock samples, created *in vitro* from standard laboratory or cultured strains of *P. falciparum* malaria (Methods, Supplementary Table 2). The second was a set of 28 DBS assessed as *P. falciparum* positive by RDT, and collected from a clinical setting in Kaoma, Zambia (Methods).

We sequenced the mock samples with the NOMADS8 panel on a Flongle Flow Cell, generating 345 thousand reads or 1.08Gbp (Fig. 2a). Of these reads, 80.0% passed the Guppy quality control filter and had identifiable sample barcodes on at least one end. We mapped these reads to the *P. falciparum* 3D7 reference genome and found that 76.2% (61.4% of the total reads) mapped successfully. To understand the causes of mapping failure, all unmapped reads were subsequently mapped to the human reference genome. Nearly all of the reads failing to map to the *P. falciparum* genome mapped successfully to the human reference (99.5%). These reads tended be shorter and of lower quality than those mapping to *P. falciparum*, and in optimisation experiments we were able to reduce them by using a higher stringency DNA size selection step after adapter ligation (Supplementary Fig. 3, Methods). The human-mapped reads remaining in this experiment (18.5% of total) were not removed by size selection, despite being shorter. Of the reads mapping to *P. falciparum*, 93.5% mapped to target regions, suggesting that `multiply` largely avoided the production of off-target amplicons. In the end, 57.5% of total sequencing reads were on-target for this experiment. A similar percentage was found to be on-target for the NOMADS8 panel when sequencing field samples using a standard MinION Flow Cell (62.1%).

**Figure 2.**
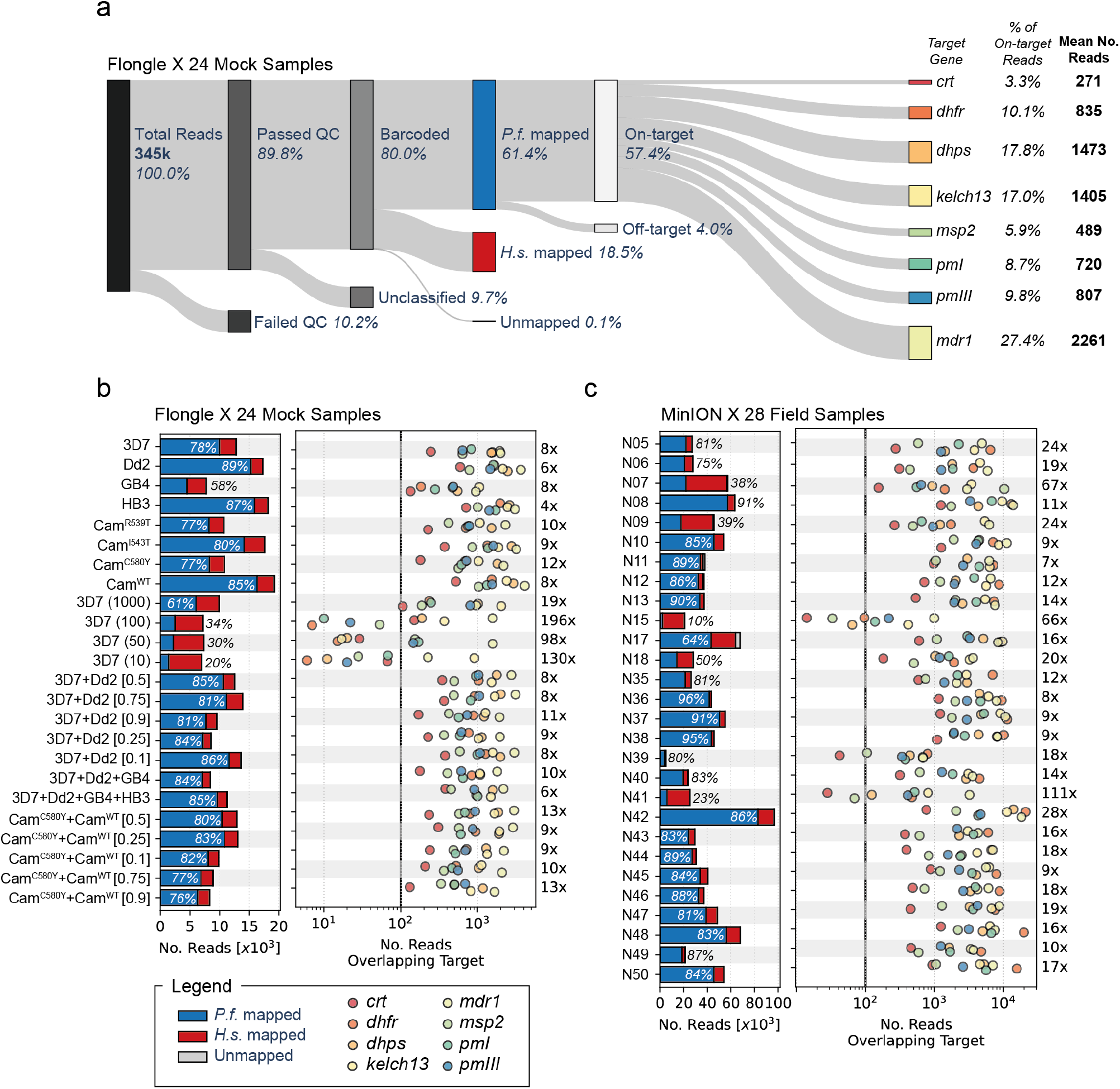
Sequencing throughput and coverage across samples and target genes for the NOMADS8 panel. (a) Diagram of reads produced on a Flongle Flow Cell (FLO-FLG001) sequencing 24 mock samples comprised of *P. falciparum* and human DNA. The leftmost bar represents all reads (n=345,000, 100%) generated during the sequencing run that are sequentially subdivided in the data analysis process to the reads of interest, i.e. those mapped to target genes (n=198,030, 57.4%). (b) Bar plot (left pane) displays the total number of reads generated for each sample stratified by mapping outcome: mapped to *P. falciparum* (P.f.) (blue), human (H.s.) (red), or failing to map (grey, too few to be visible). *P.f*. mapping percentages indicated with text. Scatter plot (right pane) displays the number of reads overlapping each target gene (labeled by colors) after mapping for each sample. Note number of reads (x-axis) is displayed in log-scale. For most samples, all target genes have >100X coverage. Number at right (e.g. 8X for 3D7) gives the fold-difference between the highest coverage and lowest coverage target. (c) Same as (b) but for 28 field samples collected as DBS from Kaoma, Zambia.

Next, we interrogated how uniformly on-target sequencing coverage was distributed across the amplicons of the NOMADS8 panel by quantifying the number of reads that overlapped each of our targets. For the mock and field samples, the median fold-difference in coverage between the highest and lowest abundance amplicons were 9.3 and 16.2, respectively (Fig. 2b,c). With both the mock and field sample sequencing runs, the rank-order of amplicons by abundance was consistent (mock samples, Spearman’s *ρ* = 0.77; field samples, Spearman’s *ρ* = 0.85; Supplementary Fig. 4a,b). This indicates that coverage variation across amplicons is largely systematic, and likely a function of differences in PCR efficiency, rather than stochastic. However, comparing mock and field sample sets, the order of amplicon abundances differed slightly, indicating sample-set dependent effects. For example, *mdr1* was lower abundance and *dhfr* was higher abundance in field samples; but notably, *mdr1* is present in multiple copies in the laboratory strain Dd2, which is used in 8 of 24 of our mock samples (Supplementary Table 2). *crt1* was the lowest abundance for both mock and field samples; also being the longest amplicon in the NOMADS8 panel (3874bp) with the second highest (A+T) composition (81.62%, behind *pmIII* with 81.66%) and most bases in long homopolymers (605bp in homopolymers length 4 or greater). Despite this, *crt1* still had a median of 230-fold coverage in the mock sample experiment and 508-fold coverage with the field samples (Supplementary Fig. 4a,b).

The NOMADS16 panel had less uniform coverage across amplicons in comparison to the NOMADS8 panel (Supplementary Fig. 5). In particular, the fold-difference between the highest and lowest abundance amplicons was 141 for the mock samples and 324 for the field samples. This was driven in part by the *hrp3* upstream amplicon producing very low median coverage relative to other amplicons in both experiments (mock samples, median of 28-fold coverage; field samples, median of 30-fold coverage; Supplementary Fig. 4c,d); with the *hrp3* upstream amplicon excluded, the fold-differences between the highest and lowest abundance amplicons was substantially reduced, but still higher than with the NOMADS8 panel (mock samples, 43.8; field samples, 51.5). As with the NOMADS8 panel, amplicons in the NOMADS16 panel had consistent relative abundances (mock samples, Spearman’s *ρ* = 0.85; field samples, Spearman’s *ρ* = 0.84 and Supplementary Fig. 4c,d), but again the specific ordered varied somewhat between mock and field samples.

### Effect of parasitemia on sequencing performance

We examined the effect that sample parasitemia had on three measures of sequencing performance: the number of reads generated per sample, normalised to the mean for the sequencing run; the percentage of those reads that mapped to *P. falciparum*; and the fold-difference in coverage between the highest and lowest abundance amplicons for the sample (Fig. 3). We evaluated these metrics for the four sequencing experiments described above, plus an additional experiment where we sequenced 96 mock samples with titrated parasitemia values, using the NOMADS16 panel on a standard MinION Flow Cell. Jointly these sample sets had parasitemia values ranging from 10 parasites per microlitre (p/*μ*L) to over 100,000p/*μ*L. Unfortunately, we note that only 12 of 28 fields samples had available parasitemia data, and all were above 1,000p/*μ*L.

**Figure 3.**
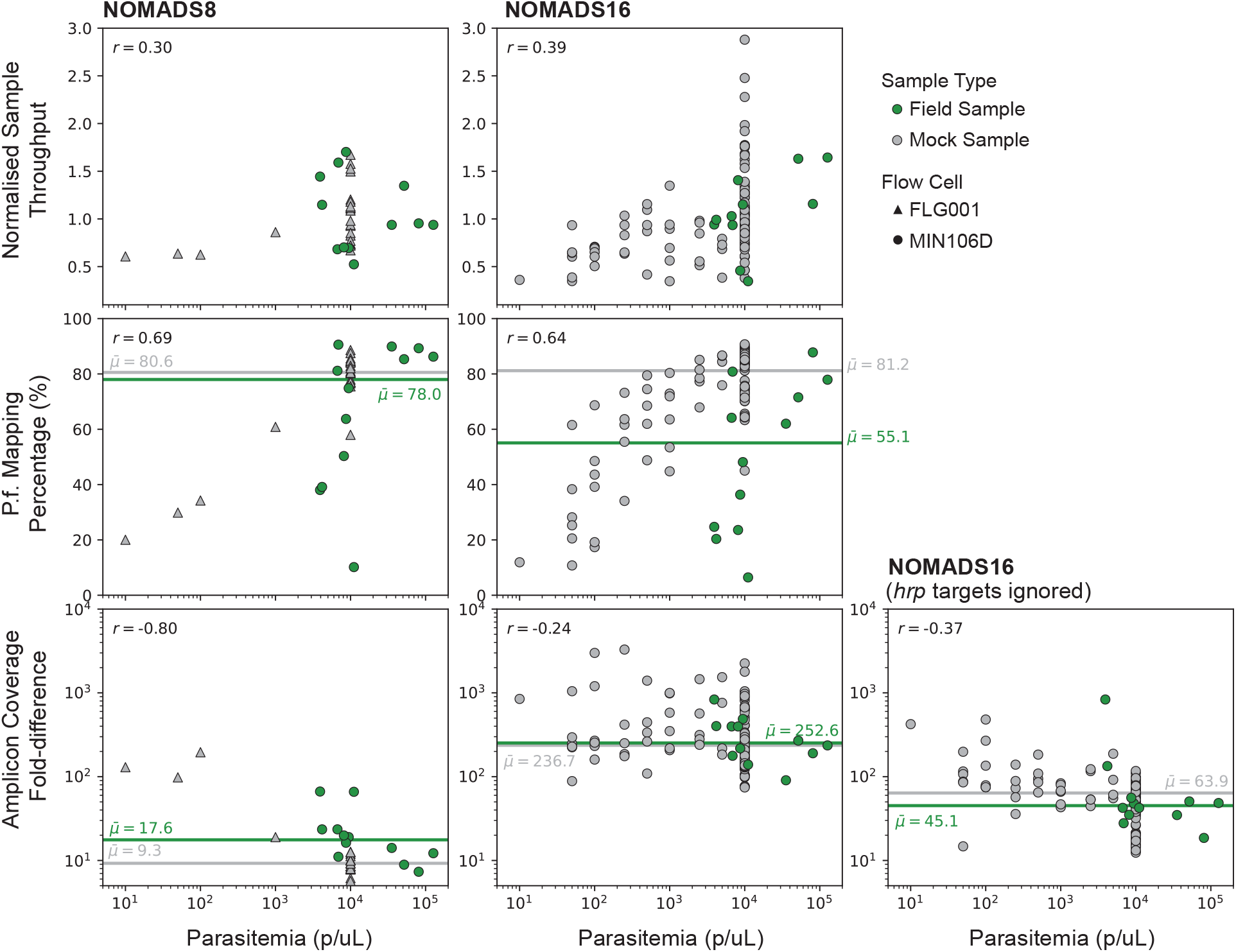
Effect of parasitemia on sequencing performance measures. Scatter plots display the effect that parasitemia (x-axis) has on the NOMADS8 (left), NOMADS16 (middle) and NOMADS16 amplicon panel with *hrp* genes ignored (right). Three measures of sequencing performance are shown (y-axis): “Normalised Sample Throughput”, which is the number of reads generated for a sample, divided by the mean number of reads per sample for the sequencing run (top row); “*P.f*. Mapping Percentage”, which is the percentage of all reads from a sample that mapped to *P. falciparum* (middle row); and the “Amplicon Coverage Fold-difference” which, for a given sample, is the ratio of the number of reads overlapping the highest abundance amplicon, divided by the number of reads overlapping the lowest abundance amplicon (bottom row). Each point is either an mock sample (grey) or a field sample (green), sequenced with a Flongle Flow Cell (FLG001, triangles) or regular MinION Flow Cell (MIN106D, circles). Median values are shown as horizontal lines for both sample types, and Pearson correlation coefficient is given in top left. Only the 12 of 28 field samples with parasitemia data are included here.

We did not perform any sample input normalisation and found that the number of reads per sample had only a weak positive correlation with parasitemia for both the NOMADS8 (Pearson’s *r* = 0.30) and NOMADS16 panels (Pearson’s *r* = 0.39). The *P. falciparum* mapping percentages had a stronger positive correlation with parasitemia (NOMADS8, Pearson’s *r* = 0.69; NOMADS16 Pearson’s *r* = 0.64); values were markedly lower below approximately 1,000p/*μ*L.

The coverage fold-difference across amplicons was higher at lower parasitemia values, producing a negative correlation that was more pronounced for the NOMADS8 (Pearson’s *r* = −0.80) than the NOMADS16 panel (Pearson’s *r* = −0.24). With the *hrp2/3* targets and their flanking genes removed, the fold-difference in coverage across the NOMADS16 panel was 5-fold less and the negative trend with parasitemia stronger (Pearson’s *r* = −0.37). In addition to the *hrp3* upstream target being low abundance, several of the titrated mock samples contained *P. falciparum* laboratory strains Dd2 and HB3, which have *hrp2* and *hrp3* deletions, respectively. This partially masked the effect of parasitemia and increased variation in coverage. For both NOMADS8 and NOMADS16 panels, roughly 1,000p/*μ*L was the threshold below which coverage variation across amplicons increased.

### SNPs are called accurately within coding sequences for clonal infections

We sought to assess how accurately molecular markers of antimalarial drug resistance could be detected with our method. Using `Clair3` to call variants^57^, we examined SNP calls for set of substitution mutations associated with drug resistance (documented by the World Health Organisation^42^) across eight of our clonal mock samples that were sequenced on a Flongle Flow Cell (Fig. 4a). For the four mock samples containing *P. falciparum* laboratory strains, we identified all expected mutations and no false positives. Similarly, for the four mock samples created from cultured *P. falciparum* strains from Cambodia with documented artemisinin resistance, we identified the expected *kelch13* mutations and no false positives.

**Figure 4.**
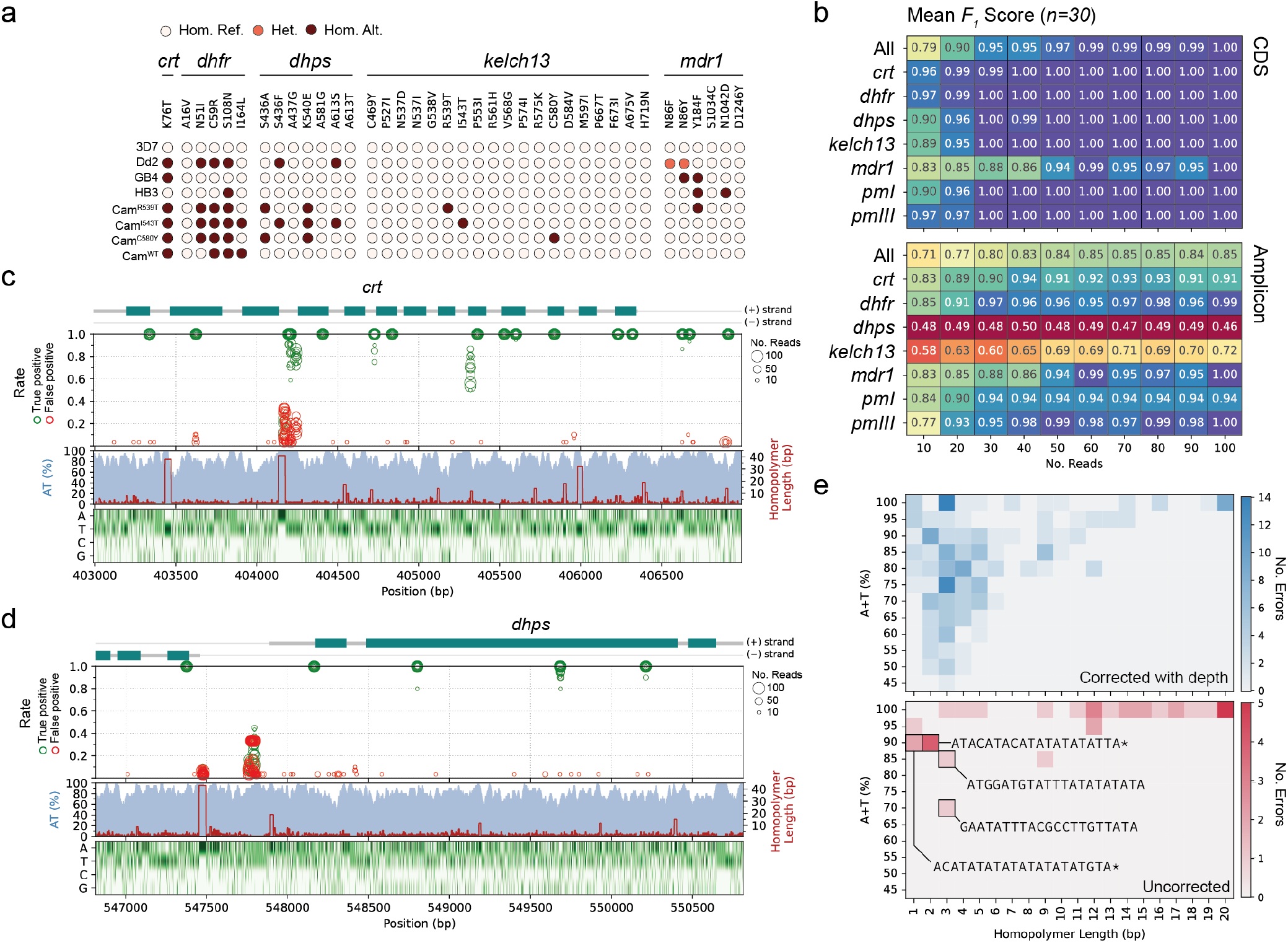
SNP calling accuracy for a set of clonal mock samples. (a) Genotype calling results from `Clair3` for eight clonal mock samples and across 38 antimalarial resistance-associated mutations. Samples were sequenced on a Flongle Flow Cell. (b) Mean *F*_1_ Score (harmonic mean of precision and recall) of SNP calling compared to PacBio data from for Dd2, GB4 and HB3 mock samples randomly downsampled to different read depths. Each square gives the mean *F*_1_ Score across thirty *in silico* replicates (ten replicates for each of Dd2, GB4 and HB4) at the indicated read depth (columns) and across the indicated region (rows). Top panel is limited to coding sequence (“CDS”) and bottom panel the entire span of the amplicons (“Amplicon”). (c) Visualisation of true positive and false positive rate of sites spanning the *crt* amplicon in chromosome 7. From top to bottom, panels show an exon diagram of *crt*; the true positive rate (green) and false positive rate (red) of each site across thirty replicates at a given read depth (indicated by circle size); A+T % in 20bp sliding windows (blue shade) and homopolymer length (red line); and heatmap of nucleotide composition. (d) Same as (c) but for *dhps* amplicon. (e) Heatmaps showing measures of sequence complexity in 20bp windows surrounding sites where errors were observed. Rows indicate A+T (%) of the 20bp window, columns indicate length of the longest hompolymer within the 20bp window, and color gives number of errors. Top panel shows errors which were corrected with additional read depth (i.e. exist at depth < 100); bottom panel shows errors that persist at a depth of 100 reads. Selected sequences are shown; * marks sequences that are an example from a bin with greater than one sequence.

Next, we expanded our analysis to examine SNP calling accuracy beyond known drug-resistance associated mutations, also characterising the effect read depth had on accuracy measures. We focused on the laboratory strains Dd2, GB4, and HB3, for which high-quality whole-genome assemblies exist^58^. Using data for each of these three mock samples, we randomly downsampled from 10 up to 100 sequencing reads, in increments of 10, for each amplicon. We repeated this procedure 10 times, to produce a total of 100 *in silico* replicates with varying read depths for each of Dd2, GB4 and HB3. For these replicates we called variants using `Clair3` and treating the whole-genome assemblies as truth, evaluated accuracy using the haplotype comparison tool `hap.py` (Methods). In Figure 4b we show the mean *F*_1_ scores (the harmonic mean of the precision and recall) for this experiment, stratified by read depth and target amplicon. The target *msp2* has been excluded as its very high sequence divergence from the reference genome makes it a case that should be handled separately, with a reference-free approach.

First we examined the coding sequences (CDS) of our targets (totalling 14.4kbp, excluding *msp2*), as these are higher complexity and are also expected to capture the overwhelming majority of possible functional mutations. At 50 reads or greater, the CDS of all targets were called with perfect accuracy except for *mdr1*. In particular, we observed 18 SNP calling errors at 5 unique sites across the 150 *in silico* replicates of *mdr1* that were downsampled to 50 reads or greater (Supplementary Table 3). Two of these SNP errors were observed in only 1 out of 150 of the *in silico* replicates, at depths of 50 and 60 reads, and both were heterozygous calls. The remaining 16 errors had two sources. First, clones of Dd2 have been observed to carry multiple copies of *mdr1*, inducing heterozygosity at codon N86 (AAT), as different copies carry N86Y (TAT) or N86F (TTT). In the complete set of reads overlapping *mdr1* in our Dd2 mock sample, the N86F mutation had a within-sample allele frequency of 58.9% (2010/3413 reads), most consistent with the mutation being carried by 1 of 2 *mdr1* copies, or 2 of 3 *mdr1* copies. In six *in silico* replicates `Clair3` called the site as homozygous N86F instead (Supplementary Fig. 6b). Second, from codons 643 to 662 in *mdr1* there is a repetitive stretch of 19 amino acids (7xAsn(AAT), 2xAsp(GAT), 10xAsn(AAT)) predicted by AlphaFold^59^ to form an unstructured loop between two transmembrane domains (Supplementary Fig. 6d). In the strain GB4, two nearby substitutions shift the position of adjacent aspartic acid residues within this region (8xAsn, 2xAsp, 9xAsn); in 20% of *in silcio* replicates at least one of these substitutions was called as heterozygous, instead of homozygous (Supplementary Fig. 6c). At 100 reads per amplicon these issues were resolved and the entire coding sequence of our panel was called perfectly across all 30 *in silico* replicates.

Finally, we expanded the analysis to the complete genomic regions spanned by our amplicons (totalling 25.2kbp, excluding *msp2*), which includes 10.8kbp of very low complexity (86% A+T) intragenic sequence. Here, at 100 reads per amplicon we observed a mean *F*_1_ score of 0.85, with considerable variation between targets. We sought to understand the causes of these SNP calling errors. First, we visualised the position of erroneous SNP calls at different read depths across our target panel (Fig. 4c, d). We observed that areas with a high false positive rate, or diminished true positive rate, tended to be in very low complexity intragenic sequence. For example, between exons 3 and 4 of *crt* there is a homopolymer of 41 A nucleotides, around which SNP errors cluster (Fig. 4c). Similarly, upstream of *dhps* there is a 50bp AT dinucleotide repeat region in which SNP errors are concentrated. To systematically evaluate the influence of sequence context on SNP calling errors, we collected all unique sites where a SNP calling error was observed in any of the 300 *in silico* replicates in our experiment. These sites were divided into two groups: those in which the SNP error could be corrected with additional read depth (*n* = 230), and those errors which remained even in replicates with 100 reads per amplicon (*n* = 35). We computed the (A+T)-content and maximum homopolymer length in a 21bp window centered on each SNP calling error (+/−10bp). SNP errors that could be corrected with additional read depth had lower (A+T)-content (mean 81.5% vs 96.1%) and shorter homopolymers in their flanking bases (mean 5.1bp vs 10.7bp) than the uncorrected SNP calls (Fig. 4e). Of the uncorrected SNP calling errors, 23/35 (66%) were situated in 21bp windows consisting of only A or T nucleotides and 19 of these contained homopolymers of length 10 or greater.

### Long-read sequencing of the surface antigen gene *msp2* provides insights into within-sample diversity

Long reads can facilitate interrogation of more complex regions of the genome. Both the NOMADS8 and NOMADS16 panel include the highly diverse surface antigen gene *msp2*, canonically used both for complexity of infection (COI) estimation and for distinguishing recrudescence from reinfection^53^. Critically, *msp2* genetic variation induces length polymorphism across a set of known repeat-containing alleles, enabling allele typing via capillary or gel electrophoresis.

We analysed reads deriving from *msp2* across our mock samples and observed length polymorphism analogous to that detected with electrophoresis approaches (Fig. 5a). To further characterise *msp2*-derived reads, we mapped them to each of the four *P. falciparum* laboratory strains used in our mock samples and labelled them by the strain to which they had the highest identity alignment (Methods). With this basic approach to allele classification, we could both confirm that the observed length polymorphism was driven by different underlying alleles of the expected types, and identify mock samples carrying multiple alleles. Next, we sought to explore an approach to read classification that avoided using *a priori* information about allele types. To this end, we implemented a global alignment algorithm for pairs of reads that used base-level quality scores to assess the likelihood that both reads derived from the same underlying haplotype sequence (Methods). Using this algorithm, we performed pairwise global alignment of all *msp2*-derived reads for each sample and hierarchically clustered the resulting pairwise alignment score matrices (Fig. 5b). In cases where a single *P. falciparum* strain was used to produce a mock sample, the pairwise alignment score matrices had little structure, consistent with a single *msp2* allele being present. In cases where multiple *P. falciparum* strains were combined to produce a mock sample, structure within the pairwise alignment matrices was consistent with multiple *msp2* alleles being present.

**Figure 5.**
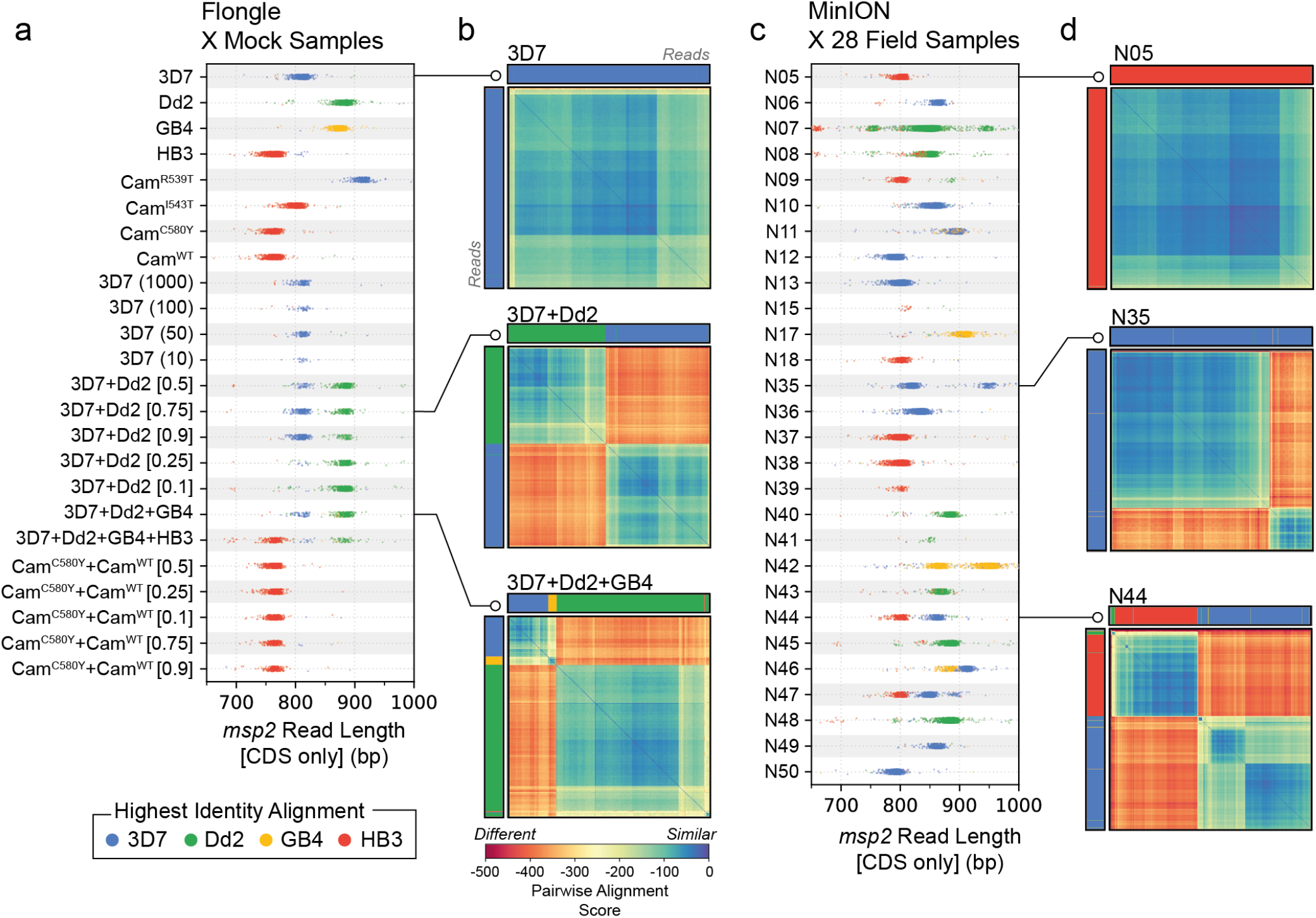
Analysis of length polymorphism and pairwise similarity of *msp2*-derived reads. (a) Read length distributions of *msp2* alleles across 24 mock samples. Each dot represents the length of a single read that was trimmed to the extent of *msp2* coding-sequence (CDS) after mapping. Individual reads are colored by the laboratory strain to which they have the highest identity alignment. Multimodal distributions are indicative of mixed infections. (b) Hierarchically clustered heatmaps of *msp2*-derived reads showing pairwise alignment scores. Each cell is colored by the global pairwise alignment score between two *msp2*-derived reads, which have been hierarchically clustered along both rows and columns. Colors of rows and columns indicate the laboratory strain to which each read has the highest identity alignment, as in panel (a). Heatmaps are shown for three different mock samples: clonal 3D7 (top); mixture of 3D7 and Dd2 (middle); and mixture of 3D7, Dd2 and GB4 (bottom). Note how reads cluster based on allele type. GB4 reads are under-represented in the bottom heatmap, likely due to lower DNA quality. (c) and (d) are the same as (a) and (b), but for 28 field samples from Zambia. In (c), read lengths distributions suggest the presence of both clonal and mixed infections. In (d), examples of likely clonal infection (top); two-strain infection (middle); and three-strain infection (bottom).

The analysis using mock samples highlighted two limitations of these approaches. First, a general limitation of using only a single locus to learn about COI is that strains within a mixed/polyclonal infection may share the same allele at that locus, leading to underestimation of COI. We observed this with mock samples of COI= 2 created by combining Cam^WT^ and Cam^C580Y^ cultured *P. falciparum* strains. Second, reads identified as deriving from GB4 were underrepresented in higher COI mock samples (Fig. 5b). This may be due to the GB4 genomic DNA we obtained being lower quality. Consistent with this, the clonal mock sample created from GB4 genomic DNA produced less reads compared with the other laboratory strains (Fig. 2b).

We applied these approaches to characterise the *msp2*-derived reads in our field sample set and observed a variety of patterns reflecting clonal and mixed infections (Fig. 5c,d).

## Discussion

Though widely deployed for the genomic surveillance of viral and bacterial pathogens, nanopore sequencing of *P. falciparum* malaria is relatively rare. Here, we have developed an approach to targeted nanopore sequencing of *P. falciparum* malaria that is flexible and cost-effective. Our approach begins with DBS as input and can produce genomic data of public health significance in 2 to 3 days at approximately USD $25 per sample. It utilises the portable MinION sequencing device, which enables sequencing in low-resource settings: including small, remote, or even mobile laboratories. Importantly, DBS collection is considerably less invasive and easier to scale than venous blood draws. Although using DBS can necessitate including sWGA as part of sample processing, we have demonstrated that its costs can be substantially reduced when combined with targeted sequencing. Moreover, in developing `multiply`, we have provided a general and principled solution to the design of multiplex PCR for targeted sequencing of *P. falciparum*. This will enable rapid creation and updating of amplicon panels as *P. falciparum*, or our knowledge of it, evolves. The software is open-source and freely available, allowing teams to design panels addressing their specific research or surveillance questions. Using `multiply`, we produced two amplicon sequencing panels containing eight- and sixteen-targets, reflecting the major public health uses of genomic data: tracking resistance to various antimalarial drugs, monitoring the sensitivity of RDTs, understanding the diversity of malaria vaccine targets and assessing within-sample diversity (which can help discriminate recrudescence from reinfection, and gives some indication of local transmission intensity). In contrast to all existing *P. falciparum* targeted sequencing approaches, our panels generate amplicons between 3 to 4kbp, thereby producing individual reads that span the entire CDS of nearly all of our target genes.

A limitation of our current method is that it has poorer performance on low parasitemia samples in comparison with other *P. falciparum* amplicon-based methods designed for short-read sequencing^22–24, 60^. Parasitemia and DBS sample quality (characterised by factors like age, storage conditions, number of blood spots and spot size) will influence the maximum amplicon length above which PCR performance will suffer due to an insufficient concentration of template DNA molecules of an adequate length. In addition, PCRs with longer amplicons typically have reduced efficiency in comparison with shorter alternatives. The 3 to 4kbp, CDS-spanning amplicons in our panel exhibited robust assay performance on mock and field samples with above ~1000 parasites per microlitre. Additional experiments, especially with field samples, are necessary to more confidently establish this threshold and determine the requirements our long-read amplicon panels put on DBS collection procedures and quality. Regardless, it is likely the NOMADS panels are best suited to higher parasitemia, symptomatic or clinical cases, rather than lower parasitemia asymptomatic cases. An advantage of developing `multiply` is that, should it be necessary, we will be able to rapidly design new amplicon panels with shorter lengths (e.g. 1-2kbp). Moreover, work on adaptive sampling of reads during nanopore sequencing has recently been applied to *P. falciparum* malaria^61^, and variations of this approach may help recover better data from low parasitemia samples.

An important question not directly addressed by this study is how sensitively our assay can detect minor clones, and the mutations they carry, in mixed/polyclonal *P. falciparum* infections^62^. The extent to which a sequencing method can detect minor clones depends on two sequential processes. The first is the reliability with which the laboratory protocol recapitulates, in the sequencing reads, the number and proportions of *P. falciparum* strains that existed in the cognate sample. This is fundamentally a sampling process, with higher variation and lower sensitivity expected in low parasitemia and low coverage samples; but it is also influenced by the non-linear dynamics of any amplification procedures that may be employed. Our approach uses sWGA, which has been shown to weaken correlations between strain proportions in a mixed infection and within-sample allele frequencies^24^. Once reads from different clones have been generated, the second process is to identify them by variant calling and/or haplotype inference, simultsneously distinguishing true variation from error or contamination. In clonal samples we determined that 30- to 50-fold coverage is sufficient for high accuracy SNP calling across all but the most extreme low complexity stretches of the genome (i.e. 100% (A+T)-content and/or hompolymers > 10bp). We highlight that our SNP accuracy analysis used data generated with the R9.4.1 Flow Cell on a Flongle, and that the new R10.4.1 Flow Cell will almost certainly exhibit better accuracy in such regions^63^. Regardless, the fact that coverage levels tens- to hundreds-of times higher than this can be readily attained gives some indication that detection of minor clones comprising ~10% of the sample may be feasible. Critically, we used `Clair3` to perform variant calling, which was designed for diploid human or haploid genomes^57^, but not for samples with varying and unknown ploidy as is the case with *P. falciparum*. In order to properly investigate the limits of minor clone detection, haplotype inference tools that can handle the polyploid nature of *P. falciparum* in conjunction with the greater length and higher error rate of nanopore reads must be developed. Going forward, these may be built by adapting existing short-read haplotype inference tools, such as `DADA2`^64^ and `SeekDeep`^65^, for nanopore data; adapting nanopore-based tools such as `Clair3`^57^ or `WhatsHap`^66^, for malaria; or be developed as new fit-to-purpose tools.

Long-read amplicon sequencing of *P. falciparum* malaria brings benefits for malaria genomic surveillance. There are three immediate examples. First, long reads, especially those spanning entire CDS, are better suited for the detection of rare and novel mutations. Approaches using smaller reads typically focus on ~250bp regions around known, common mutations, and have primer binding locations within the target gene. Therefore, novel mutations can emerge undetected or disrupt primer annealing, ultimately requiring the redesign or expansion of an amplicon panel. For *P. falciparum* a critical surveillance region is the propeller domain of *kelch13*^67^, which harbors an expanding list of mutations conferring artemisinin resis-tance^4, 42^, but at 855bp is too long to capture with a single amplicon in short-read sequencing. Second, once suitable computational tools are developed, long reads will enable epidemiologically relevent read-based phasing of variants within target genes^66^. For example, pyrimethamine treatment failure is predicted on the basis of a triple mutation within *dhfr* including N51I, C59R and S108N^48^; however, in some geographies single, double, and triple mutations exist, complicating this prediction for mixed infections^49, 68, 69^. Third, longer reads allow for better mapping in structurally complex or repetitive regions of the genome, and can assist with structural variant detection^32^. The investigation of several control-relevant regions of *P. falciparum*, including *msp2*, the histidine-rich proteins *hrp2* and *hrp3*, and the vaccine target *csp*; will all benefit from long-read sequencing.

## Methods

### Development of multiplex PCR panels

The NOMADS8 and NOMADS16 panels were generated using a beta version of `multiply`, called `pf-multiply`, which is available at https://github.com/JasonAHendry/pf-multiply (design file for NOMADS8; design file for NOMADS16). Both use a BED (*.bed) file to delineate the *mdr1* amplicon. NOMADS8 was generated first using the command: `python multiply.py -d designs/pf-nomads8-mdr1part.ini` NOMADS16 was created by using the augment command of `pf-multiply`. The NOMADS8 multiplex PCR primers were combined at equimolar amounts (10uM); after an initial sequencing run with mock samples on a Flongle Flow Cell, primer concentrations were crudely adjusted (doubled or halved) based on observed amplicon abundances. The same procedure was repeated with NOMADS16; i.e. one round of primer concentration adjustment was performed. Final primer sequences and concentrations are available in the protocol on protocols.io.

### Creating mock samples of *P. falciparum* and human DNA

We ordered *Plasmodium falciparum* genomic DNA for laboratory strains 3D7, Dd2, GB4 and HB3 and Cambodian field derived strains IPC 5202 (*kelch13* R539T); IPC 4912 (*kelch13* I543T), IPC 3445 (*kelch13* C580Y); and IPC 3663 (*kelch13* WT)^67^ from BEI resources (www.beiresources.org). To create 10,000p/*μ*l *in vitro* DNA mixtures we diluted these stocks to 0.25ng/*μ*l using 25ng/*μ*L human genomic DNA that was pooled from 36 HapMap cell lines^70^. DNA mixtures were then combined at different numbers and ratios to replicate mixed infections of different proportions or complexity of infection (COI), and/or serial diluted in additional human genomic DNA to replicate lower parasitemia infections (Supplementary Table 2).

### Collection of field samples

Samples were collected under an ethical waiver granted by the National Health Research Authority, Zambia under the “Laboratory Quality Improvement Research In Ministry of Health Laboratories” (NHRA000004/16/11/2021). Symptomatic patients visiting a clinic in Kaoma, Western Province, Zambia were diagnosed with an RDT while a microscopy slide and DBS (on Whatmann No3 filter paper) were also collected. All samples were deidentified and no metadata was recorded for any patient Bulk DNA was extracted from DBS using a Qiagen QIAamp Kit following manufacturers instructions. Parasitemia was quantified by light microscopy from thin film blood slides.

### Laboratory protocol and sequencing

The complete laboratory protocol is available on protocols.io as ‘Cost-effective targeted nanopore sequencing of P. falciparum malaria’.

### Bioinformatics pipeline

FAST5 files generated by the `MinKNOW` software were basecalled using Guppy 5.0.11 with a minimum quality score threshold of 8. For the Flongle experiment we used the super-accurate basecalling model and for all other experiments we used high accuracy basecalling model. FASTQ files were then demultiplexed using the same version of Guppy, without setting the `--require_both_ends` flag; e.g. with single-end barcoding. Demultiplexed FASTQ files were mapped to version 52 of the *Plasmodium falciparum* 3D7 reference genome downloaded from [plasmodb](https://plasmodb.org/common/downloads/release-52/Pfalciparum3D7/) using `minimap2` version 2.24-r1122 and the `-ax-ont` parameter setting. In the resultant BAM (*.bam) file, reads failing to map to the 3D7 reference were identified using the command `samtools view -f 0×904`, and were converted back to FASTQ files using `samtools fastq` before being remapped to the GRCh38 human reference genome, downloaded from [NCBI website](https://www.ncbi.nlm.nih.gov/assembly/GCF_000001405.26/). Reads deriving from targets were defined as those that overlap the coding-sequence determined from the Gene Feature Format (GFF) (*.gff), which was also downloaded [plasmodb](https://plasmodb.org/common/downloads/release-52/Pfalciparum3D7/gff/data/). Variant calling of reads mapping to the 3D7 reference genome was performed using the using the singularity image of `Clair3` v0.1-r12 (https://github.com/HKU-BAL/Clair3) in diploid mode with the flags `--platform=’ont’ --include_all_ctgs --enable_phasing` set.

### SNP calling accuracy analysis

#### Downsampling reads

We partitioned reads mapped to the 3D7 reference into those overlapping each of our target genes using `samtools` version 1.16, thereby producing BAM files for each of our targets. For each target BAM, we downsampled reads using the “view” command and “-s/–subsample” flag to achieve the desired number of reads. This procedure was repeated for the Dd2, GB4, and HB4 samples; downsampling to 10, 20, 30, 40, 50, 60, 70, 80, 90 and 100 reads for each target, and ten times for each number of reads. As a result, for each given depth and target, we produced 10 randomly downsampled BAM files for Dd2, GB4 and Hb3; or 30 replicates total. In Figure 4b the “All” category was created be concatenating the BAM files generated in this way for all targets in the NOMADS8 panel, excepting *msp2*. *Creating a set of true variants*. Dd2, GB4, and HB3 have been sequenced to high depth on the Pacific Bioscience Sequencing SMRT technology and the FASTA sequences from their assembly have been deposited on [plasmodb](https://plasmodb.org/common/downloads/release-52/). To identify variants in these assemblies with respect to the 3D7 reference genome, we simulated error-free reads *in silico* from the FASTA files, mapped them to the 3D7 reference genome with minimap2, and then identified variants using the `bcftools` v1.16 “mpileup” and “call” commands. In particular, we simulated 60 error-free reads for each target in our NOMADS8 panel by extracting the FASTA sequence spanning +/−4kbp of the target, based on GFF files for Dd2, GB4, and HB4 that were also downloaded from plasmodb. We arbitrarily gave all bases of these *in silico* reads a Phred quality score of 60 and reverse-complimented half of them to represent negative strand-derived reads. We then generated FASTQ files from these reads before the variant calling with `bcftools`.

#### Stratified variant call comparisons

We used the tool `hap.py` (https://github.com/Illumina/hap.py) which we downloaded as a `Docker` image (jmcdani20/hap.py:v0.3.12) to compute measures of variant calling accuracy across different target regions in comparison to the true variant set described above. To restrict accuracy measure analysis to coding sequence, we subset the 3D7 GFF downloaded from plasmodb to only “CDS” features, identified the rows pertaining to our targets, and output the chromosome, start, and stop positions as a BED (*.bed) file. We used then used the `--stratifications` flag of `hap.py` to compute measures over these intervals. We used the annotated VCF (*.vcf) files produced by hap.py to generate positional plots of false- and true-positive rate across targets.

### Analysis of *msp2* reads

#### Computing coding sequence lengths

After mapping reads to the 3D7 reference genome with `minimap2`, we extracted reads that completely overlapped the *msp2* (PF3D7_0206800) coding sequence using `bedtools intersect -F 1.0`. From the resultant BAM file, we trimmed these reads to the extent of the *msp2* coding sequencing by keeping only the section of each read that aligned within the interval [273689, 274507] of chromosome 2 (Pf3D7_02_v3); indels were retained if the bases on either side of them aligned within the interval. Unusually short trimmed reads (<400bp) were removed as likely artefacts. Trimmed reads were used to create length distribution plots. They were independently mapped, using `minimap2`, to the reference genomes for 3D7, Dd2, GB4, and HB3 (downloaded from [plasmodb](https://plasmodb.org/common/downloads/release-52/)). We let `minimap2` output a PAF (*.paf) file and computed the identity of the mapping alignment by dividing column 10 (number of matches in alignment) by column 11 (total alignment length).

#### Global pairwise alignment

We implemented a banded version of the Needleman-Wunsch algorithm to compute global alignment scores between pairs of trimmed *msp2* reads. We parameterised the scoring model such that scores reflect the log-probability that the two observed reads derived from the same underlying haplotype sequence; i.e. that all alignment differences were caused by sequencing error. Assuming an indel rate of 5%, which is broadly consistent with empirical observation, we used a linear gap score of *log*_10_(0.05). For substitution scores, we took into account the base quality scores generated by Guppy as follows. Defining *x* and *y* as the two observed bases in the match, the likelihood that they were generated from the same haplotype base *h* is

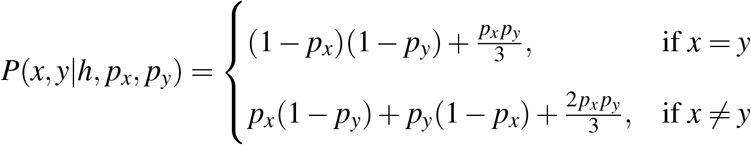

where 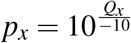 and 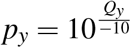, with *Q_x_* and *Q_y_* being the Phred-scaled base quality scores for *x* and *y*. The substitution score is then computed as *log*_10_(*P*(*x, y*|*h, p_x_, p_y_*)). For all alignments in this study, a band width of 80bp centered on the diagonal of the global alignment matrix was used. Hierarchical clustering of the resulting scores was performed using the `scipy.cluster.hierachy.linkage` function from the SciPy library.

## Code availability

`multiply` is available at https://github.com/JasonAHendry/multiply. Bioinformatics pipeline, SNP calling accuracy, and *msp2* analysis code is available at https://github.com/JasonAHendry/nomadic2.

## Acknowledgements

The NOMADS project is funded by the Bill & Melinda Gates Foundation (INV-003660, INV-048316). The research was supported by the Wellcome Trust Core Award Grant Number 203141/Z/16/Z with additional support from the NIHR Oxford BRC. The views expressed are those of the author(s) and not necessarily those of the NHS, the NIHR or the Department of Health. We are grateful to all health workers and patients who supported collection of field samples in Zambia. We thank Kirk Rockett for providing laboratory space and support during initial stages of the project; staff at the Oxford Genomics Centre for sequencing support, especially Amy Trebes and David Buck; Nada Kubikova for discussions about multiplex PCR primer design; Gavin Band and Annie Forster for beta testing of `multiply`; Robert Verity and Shazia Ruybal for discussions about *msp2* analysis. We also acknowledge the support of the Royal Geographical Society with IBG and Jaguar Land Rover for instigating initial collaborations through the 2018 RGS Land Rover Bursary which was awarded to J.A.H, I.G and G.B.B.

## Author contributions statement

M.C.: Methodology, Investigation, Writing - Review & Editing. M.M.: Methodology, Investigation, Writing - Review & Editing. A.J.: Methodology, Investigation, Validation, Writing - Review & Editing. J.C.: Resources; C.D.: Project administration, Writing - Review & Editing. K.S.: Project administration, Funding acquisition, Writing - Review & Editing. I.G.: Conceptualization, Funding acquisition, Writing - Review & Editing. G.B.: Conceptualization, Funding acquisition, Writing - Review & Editing. B.H.: Project administration, Funding acquisition. M.H.: Project administration, Resources. D.J.B.: Conceptualization, Supervision, Project administration, Funding acquisition, Writing - Review & Editing. J.A.H.: Conceptualization, Supervision, Funding acquisition, Investigation, Formal analysis, Software, Visualization, Writing - Original Draft.

## References

1. White, N. J. Antimalarial drug resistance. The J. clinical investigation 113, 1084–1092, DOI: 10.1172/JCI21682 (2004).

2. Haldar, K., Bhattacharjee, S. & Safeukui, I. Drug resistance in plasmodium. Nat. Rev. Microbiol. 16, 156–170, DOI: 10.1038/nrmicro.2017.161 (2018).

3. (WHO), W. H. O. World malaria report 2022. Report, WHO (2022).

4. Genomic epidemiology of artemisinin resistant malaria. Elife 5, DOI: 10.7554/eLife.08714 (2016).

5. Imwong, M. et al. The spread of artemisinin-resistant plasmodium falciparum in the greater mekong subregion: a molecular epidemiology observational study. Lancet Infect Dis 17, 491–497, DOI: 10.1016/s1473-3099(17)30048-8 (2017).

6. Hamilton, W. L. et al. Evolution and expansion of multidrug-resistant malaria in southeast asia: a genomic epidemiology study. Lancet Infect Dis 19, 943–951, DOI: 10.1016/s1473-3099(19)30392-5 (2019).

7. Balikagala, B. et al. Evidence of Artemisinin-Resistant Malaria in Africa. New Engl. J. Medicine 385, 1163–1171, DOI: 10.1056/NEJMoa2101746 (2021).

8. Uwimana, A. et al. Emergence and clonal expansion of in vitro artemisinin-resistant Plasmodium falciparum kelch13 R561H mutant parasites in Rwanda. Nat. medicine DOI: 10.1038/s41591-020-1005-2 (2020). 32747827.

9. Amoah, L. E., Abankwa, J. & Oppong, A. Plasmodium falciparum histidine rich protein-2 diversity and the implications for PfHRP 2: Based malaria rapid diagnostic tests in Ghana. Malar. J. 15, 101, DOI: 10.1186/s12936-016-1159-z (2016). 26891848.

10. Menegon, M. et al. Identification of Plasmodium falciparum isolates lacking histidine-rich protein 2 and 3 in Eritrea. Infect. Genet. Evol. J. Mol. Epidemiol. Evol. Genet. Infect. Dis. 55, 131–134, DOI: 10.1016/j.meegid.2017.09.004 (2017). 28889944.

11. Berhane, A. et al. Major Threat to Malaria Control Programs by Plasmodium falciparum Lacking Histidine-Rich Protein 2, Eritrea. Emerg. Infect. Dis. 24, 462–470, DOI: 10.3201/eid2403.171723 (2018). 29460730.

12. Golassa, L., Messele, A., Amambua-Ngwa, A. & Swedberg, G. High prevalence and extended deletions in Plasmodium falciparum hrp2/3 genomic loci in Ethiopia. PloS One 15, e0241807 (2020). 33152025.

13. Feleke, S. M. et al. Plasmodium falciparum is evolving to escape malaria rapid diagnostic tests in Ethiopia. Nat. Microbiol. 6, 1289–1299, DOI: 10.1038/s41564-021-00962-4 (2021). 34580442.

14. Ariey, F. et al. A molecular marker of artemisinin-resistant plasmodium falciparum malaria. Nature 505, 50–5, DOI: 10.1038/nature12876 (2014).

15. Miotto, O. et al. Genetic architecture of artemisinin-resistant plasmodium falciparum. Nat. Genet. 47, 226–234, DOI: 10.1038/ng.3189 (2015).

16. Gamboa, D. et al. A large proportion of p. falciparum isolates in the amazon region of peru lack pfhrp2 and pfhrp3: Implications for malaria rapid diagnostic tests. PLOS ONE 5, e8091, DOI: 10.1371/journal.pone.0008091 (2010).

17. Cheng, Q. et al. Plasmodium falciparum parasites lacking histidine-rich protein 2 and 3: a review and recommendations for accurate reporting. Malar. J. 13, 283, DOI: 10.1186/1475-2875-13-283 (2014).

18. Gardner, M. J. et al. Genome sequence of the human malaria parasite plasmodium falciparum. Nature 419, 498–511, DOI: 10.1038/nature01097 (2002).

19. Rodríguez-Gijón, A. et al. A genomic perspective across earth’s microbiomes reveals that genome size in archaea and bacteria is linked to ecosystem type and trophic strategy. Front Microbiol 12, 761869, DOI: 10.3389/fmicb.2021.761869 (2021).

20. Martinez-Gutierrez, C. A. & Aylward, F. O. Genome size distributions in bacteria and archaea are strongly linked to evolutionary history at broad phylogenetic scales. PLoS Genet. 18, e1010220, DOI: 10.1371/journal.pgen.1010220 (2022).

21. Cui, J., Schlub, T. E. & Holmes, E. C. An allometric relationship between the genome length and virion volume of viruses. J. Virol. 88, 6403–6410, DOI: doi:10.1128/JVI.00362-14 (2014).

22. Jacob, C. G. et al. Genetic surveillance in the greater mekong subregion and south asia to support malaria control and elimination. Elife 10, DOI: 10.7554/eLife.62997 (2021).

23. Tessema, S. K. et al. Sensitive, highly multiplexed sequencing of microhaplotypes from the plasmodium falciparum heterozygome. J Infect Dis 225, 1227–1237, DOI: 10.1093/infdis/jiaa527 (2022).

24. LaVerriere, E. et al. Design and implementation of multiplexed amplicon sequencing panels to serve genomic epidemiology of infectious disease: A malaria case study. Mol Ecol Resour 22, 2285–2303, DOI: 10.1111/1755-0998.13622 (2022).

25. Aydemir, O. et al. Drug-resistance and population structure of plasmodium falciparum across the democratic republic of congo using high-throughput molecular inversion probes. J Infect Dis 218, 946–955, DOI: 10.1093/infdis/jiy223 (2018).

26. Verity, R. et al. The impact of antimalarial resistance on the genetic structure of plasmodium falciparum in the drc. Nat. Commun. 11, 2107, DOI: 10.1038/s41467-020-15779-8 (2020).

27. Tegally, H. et al. The evolving sars-cov-2 epidemic in africa: Insights from rapidly expanding genomic surveillance. Science 378, eabq5358, DOI: 10.1126/science.abq5358 (2022).

28. Loman, N. J. & Watson, M. Successful test launch for nanopore sequencing. Nat Methods 12, 303–4, DOI: 10.1038/nmeth.3327 (2015).

29. Quick, J. et al. Real-time, portable genome sequencing for ebola surveillance. Nature 530, 228–232, DOI: 10.1038/nature16996 (2016).

30. Faria, N. R. et al. Establishment and cryptic transmission of zika virus in brazil and the americas. Nature 546, 406–410, DOI: 10.1038/nature22401 (2017).

31. Payne, A., Holmes, N., Rakyan, V. & Loose, M. Bulkvis: a graphical viewer for oxford nanopore bulk fast5 files. Bioinformatics 35, 2193–2198, DOI: 10.1093/bioinformatics/bty841 (2019).

32. Amarasinghe, S. L. et al. Opportunities and challenges in long-read sequencing data analysis. Genome Biol. 21, 30, DOI: 10.1186/s13059-020-1935-5 (2020).

33. Runtuwene, L. R. et al. Nanopore sequencing of drug-resistance-associated genes in malaria parasites, Plasmodium falciparum. Sci. reports 8, 8286, DOI: 10.1038/s41598-018-26334-3 (2018). 29844487.

34. Razook, Z. et al. Real time, field-deployable whole genome sequencing of malaria parasites using nanopore technology. bioRxiv DOI: 10.1101/2020.12.17.423341 (2020). https://www.biorxiv.org/content/early/2020/12/18/2020.12.17.423341.full.pdf.

35. White, M. T. et al. Immunogenicity of the rts,s/as01 malaria vaccine and implications for duration of vaccine efficacy: secondary analysis of data from a phase 3 randomised controlled trial. The Lancet Infect. Dis. 15, 1450–1458, DOI: 10.1016/S1473-3099(15)00239-X (2015).

36. Datoo, M. S. et al. Efficacy of a low-dose candidate malaria vaccine, r21 in adjuvant matrix-m, with seasonal administration to children in burkina faso: a randomised controlled trial. The Lancet 397, 1809–1818, DOI: 10.1016/S0140-6736(21)00943-0 (2021).

37. Quick, J. et al. Multiplex pcr method for minion and illumina sequencing of zika and other virus genomes directly from clinical samples. Nat Protoc 12, 1261–1276, DOI: 10.1038/nprot.2017.066 (2017).

38. Untergasser, A. et al. Primer3plus, an enhanced web interface to primer3. Nucleic Acids Res 35, W71–4, DOI: 10.1093/nar/gkm306 (2007).

39. Johnston, A. D., Lu, J., Ru, K. L., Korbie, D. & Trau, M. Primerroc: accurate condition-independent dimer prediction using roc analysis. Sci Rep 9, 209, DOI: 10.1038/s41598-018-36612-9 (2019).

40. Altschul, S. F., Gish, W., Miller, W., Myers, E. W. & Lipman, D. J. Basic local alignment search tool. J Mol Biol 215, 403–10, DOI: 10.1016/s0022-2836(05)80360-2 (1990).

41. Camacho, C. et al. Blast+: architecture and applications. BMC Bioinforma. 10, 421, DOI: 10.1186/1471-2105-10-421 (2009).

42. (WHO), W. H. O. Report on antimalarial drug efficacy, resistance and response: 10 years of surveillance (2010-2019). Report, WHO (2020).

43. Su, X. Z., Wu, Y., Sifri, C. D. & Wellems, T. E. Reduced extension temperatures required for pcr amplification of extremely a+t-rich dna. Nucleic Acids Res 24, 1574–5, DOI: 10.1093/nar/24.8.1574 (1996).

44. Miller, R. H. et al. A deep sequencing approach to estimate plasmodium falciparum complexity of infection (coi) and explore apical membrane antigen 1 diversity. Malar J 16, 490, DOI: 10.1186/s12936-017-2137-9 (2017).

45. Fidock, D. A. et al. Mutations in the p. falciparum digestive vacuole transmembrane protein pfcrt and evidence for their role in chloroquine resistance. Mol Cell 6, 861–71, DOI: 10.1016/s1097-2765(05)00077-8 (2000).

46. Djimdé, A., Doumbo, O. K., Steketee, R. W. & Plowe, C. V. Application of a molecular marker for surveillance of chloroquine-resistant falciparum malaria. The Lancet 358, 890–891, DOI: 10.1016/S0140-6736(01)06040-8 (2001).

47. Cowman, A. F., Morry, M. J., Biggs, B. A., Cross, G. A. & Foote, S. J. Amino acid changes linked to pyrimethamine resistance in the dihydrofolate reductase-thymidylate synthase gene of plasmodium falciparum. Proc Natl Acad Sci U S A 85, 9109–13, DOI: 10.1073/pnas.85.23.9109 (1988).

48. Plowe, C. V. et al. Mutations in plasmodium falciparum dihydrofolate reductase and dihydropteroate synthase and epidemiologic patterns of pyrimethamine-sulfadoxine use and resistance. J Infect Dis 176, 1590–6, DOI: 10.1086/514159 (1997).

49. Roper, C. et al. Intercontinental spread of pyrimethamine-resistant malaria. Science 305, 1124, DOI: 10.1126/science.1098876 (2004).

50. Brooks, D. R. et al. Sequence variation of the hydroxymethyldihydropterin pyrophosphokinase: dihydropteroate synthase gene in lines of the human malaria parasite, plasmodium falciparum, with differing resistance to sulfadoxine. Eur J Biochem. 224, 397–405, DOI: 10.1111/j.1432-1033.1994.00397.x (1994).

51. Triglia, T., Wang, P., Sims, P. F., Hyde, J. E. & Cowman, A. F. Allelic exchange at the endogenous genomic locus in plasmodium falciparum proves the role of dihydropteroate synthase in sulfadoxine-resistant malaria. The EMBO J. 17, 3807–3815, DOI: https://doi.org/10.1093/emboj/17.14.3807 (1998).

52. Price, R. N. et al. Mefloquine resistance in plasmodium falciparum and increased pfmdr1 gene copy number. Lancet 364, 438–447, DOI: 10.1016/s0140-6736(04)16767-6 (2004).

53. Snounou, G. & Beck, H. P. The use of pcr genotyping in the assessment of recrudescence or reinfection after antimalarial drug treatment. Parasitol Today 14, 462–7, DOI: 10.1016/s0169-4758(98)01340-4 (1998).

54. Amato, R. et al. Genetic markers associated with dihydroartemisininx2013;piperaquine failure in *plasmodium falciparum* malaria in cambodia: a genotypex2013;phenotype association study. The Lancet Infect. Dis. 17, 164–173, DOI: 10.1016/S1473-3099(16)30409-1 (2017).

55. Witkowski, B. et al. A surrogate marker of piperaquine-resistant plasmodium falciparum malaria: a phenotype-genotype association study. The Lancet Infect. Dis. 17, 174–183, DOI: 10.1016/S1473-3099(16)30415-7 (2017).

56. Josh, Q. One-pot ligation protocol for oxford nanopore libraries. protocols.io (2018). https://dx.doi.org/10.17504/protocols.io.k9acz2e.

57. Zheng, Z. et al. Symphonizing pileup and full-alignment for deep learning-based long-read variant calling. Nat. Comput. Sci. 2, 797–803, DOI: 10.1038/s43588-022-00387-x (2022).

58. Otto, T. D. et al. Long read assemblies of geographically dispersed plasmodium falciparum isolates reveal highly structured subtelomeres. Wellcome Open Res 3, 52, DOI: 10.12688/wellcomeopenres.14571.1 (2018).

59. Jumper, J. et al. Highly accurate protein structure prediction with alphafold. Nature 596, 583–589, DOI: 10.1038/s41586-021-03819-2 (2021).

60. Early, A. M. et al. Detection of low-density plasmodium falciparum infections using amplicon deep sequencing. Malar J 18, 219, DOI: 10.1186/s12936-019-2856-1 (2019).

61. De Meulenaere, K., Cuypers, W. L., Rosanas-Urgell, A., Laukens, K. & Cuypers, B. Selective whole-genome sequencing of *plasmodium* parasites directly from blood samples by nanopore adaptive sampling. bioRxiv 2022.11.29.518068, DOI: 10.1101/2022.11.29.518068 (2022).

62. Lerch, A. et al. Development of amplicon deep sequencing markers and data analysis pipeline for genotyping multi-clonal malaria infections. BMC Genomics 18, 864, DOI: 10.1186/s12864-017-4260-y (2017).

63. Sereika, M. et al. Oxford nanopore r10.4 long-read sequencing enables the generation of near-finished bacterial genomes from pure cultures and metagenomes without short-read or reference polishing. Nat Methods 19, 823–826, DOI: 10.1038/s41592-022-01539-7 (2022).

64. Callahan, B. J. et al. Dada2: High-resolution sample inference from illumina amplicon data. Nat Methods 13, 581–3, DOI: 10.1038/nmeth.3869 (2016).

65. Hathaway, N. J., Parobek, C. M., Juliano, J. J. & Bailey, J. A. Seekdeep: single-base resolution de novo clustering for amplicon deep sequencing. Nucleic Acids Res 46, e21, DOI: 10.1093/nar/gkx1201 (2018).

66. Martin, M. et al. Whatshap: fast and accurate read-based phasing, DOI: 10.1101/085050 (2016).

67. Straimer, J. et al. Drug resistance. k13-propeller mutations confer artemisinin resistance in plasmodium falciparum clinical isolates. Science 347, 428–31, DOI: 10.1126/science.1260867 (2015).

68. Roper, C. et al. Antifolate antimalarial resistance in southeast africa: a population-based analysis. Lancet 361, 1174–81, DOI: 10.1016/s0140-6736(03)12951-0 (2003).

69. null, n. et al. Pf7: an open dataset of plasmodium falciparum genome variation in 20,000 worldwide samples [version 1; peer review: awaiting peer review]. Wellcome Open Res. 8, DOI: 10.12688/wellcomeopenres.18681.1 (2023).

70. The international hapmap project. Nature 426, 789–96, DOI: 10.1038/nature02168 (2003).

