## Supplementary Materials for "Flexible and cost-effective genomic surveillance of *P. falciparum* malaria with targeted nanopore sequencing"

February 6, 2023

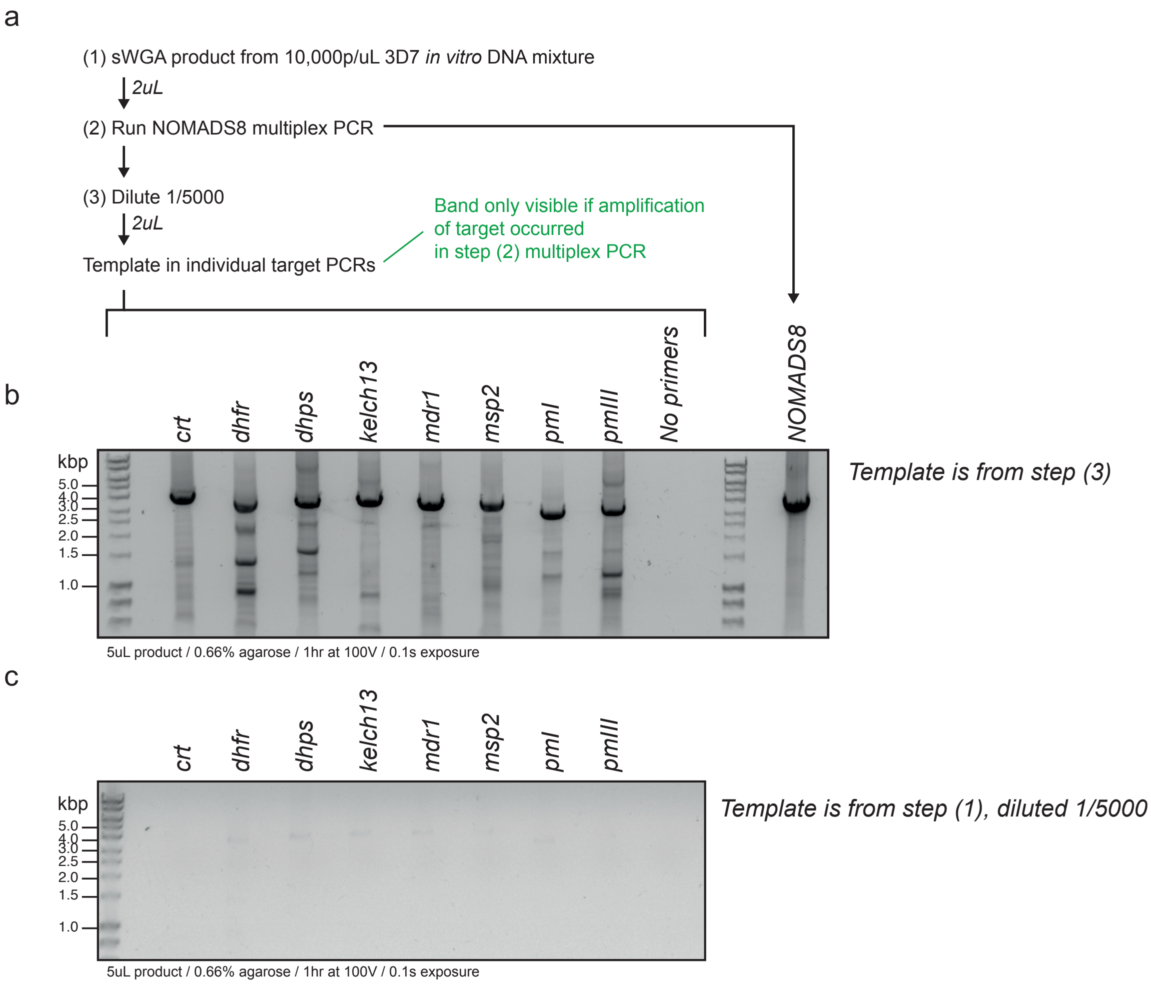


**Supplementary Figure 1: Workflow for confirmation of NOMADS8 target amplification by agarose gel electrophoresis**. (a) Diagram of workflow. sWGA is performed on mock sample made from the 3D7 reference strain (Methods). NOMADS8 multiplex PCR is performed on 2$\mu$L of sWGA product; product is shown in rightmost lane of (b). Multiplex PCR product is then diluted 1/5,000 before being split into 2$\mu$L aliquots which are used as template DNA in individual target PCRs. (b) Product from individual PCRs. Only targets with successful amplification in the multiplex PCR will have sufficient template for a successful individual PCR. (c) Negative control showing product of individual PCRs when a 1/5,000 dilution of the sWGA product is used, i.e. in the absence of the multiplex PCR. Difference between (b) and (c) validates that multiplex PCR is necessary for successful individual PCRs.

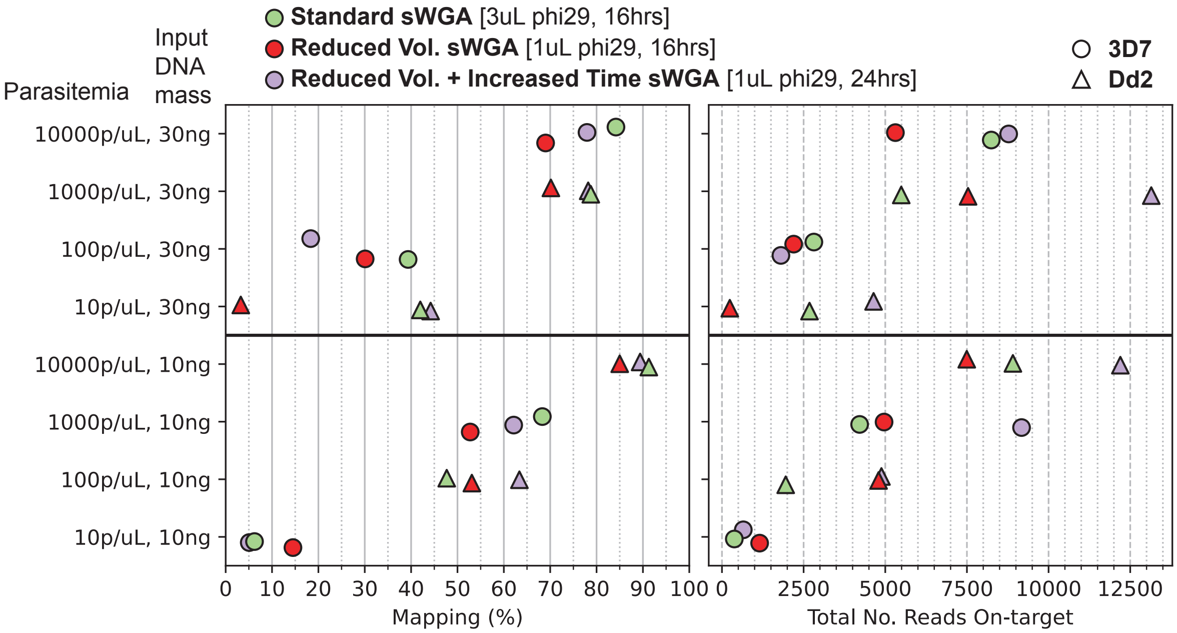


**Supplementary Figure 2: A reduced sWGA reaction volume allows for adequate coverage of target amplicons.** Three sWGA reaction conditions (marker color) were tested across four parasitemia levels (y-axis) and two input DNA mass amounts (top and bottom panel) using mock samples created from two different lab strains of *P. falciparum* (marker shape). Sequencing was completed on a Flongle Flow Cell using the NOMADS8 panel. *P. falciparum* mapping percentage (left pane) and the number of reads overlapping target genes (right pane) are shown. Reduced volume sWGA does consistently decrease *P. falciparum* mapping percentage, but still allows for robust coverage over target genes.

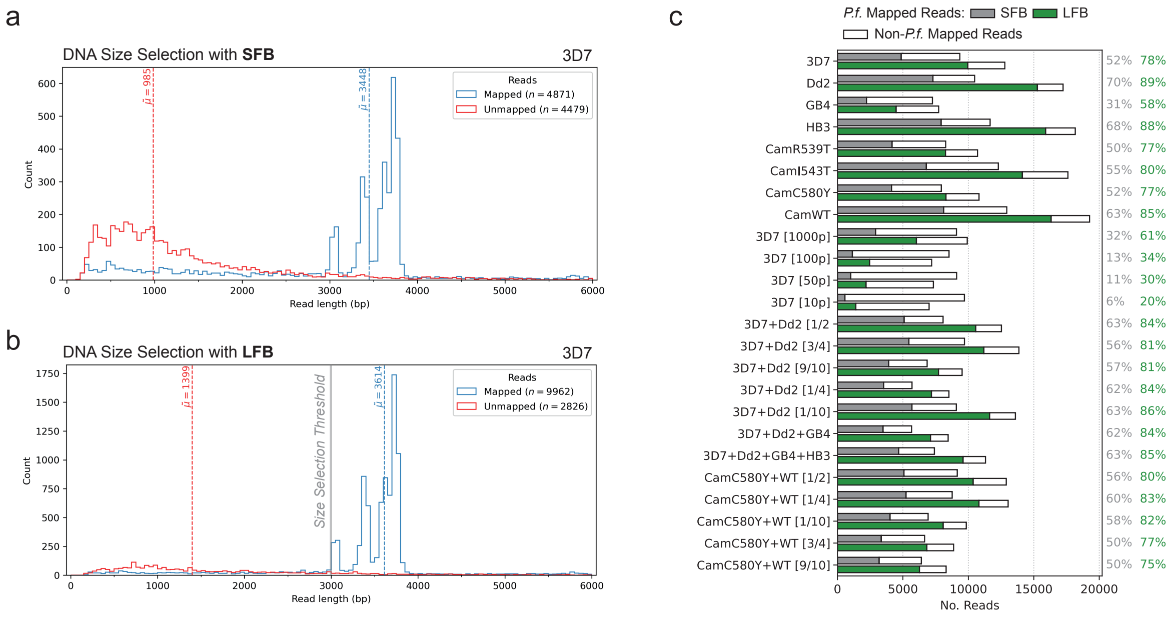


**Supplementary Figure 3: Effect of increased stringency DNA size selection on *P. falciparum* mapping percentages with the NOMADS8 panel**. Oxford Nanopore Technologies provides the option of two different DNA size selection buffers after adapter ligation and prior to sequencing: 'SFB', for short-fragment buffer, which recovers all DNA fragments; 'LFB', for long-fragment buffer, which selects for fragments $>$ 3kbp. (a) Read length distribution for mock 3D7 sample at 10,000p/$\mu$L, using SFB size selection. Blue shows reads mapped to *P. falciparum* 3D7 reference; red shows all unmapped reads. (b) Exact same PCR products sequenced in panel (a), but using LFB size selection during library preparation. Short, unmapped reads are dramatically reduced. (c) Barplot showing total reads for mock samples sequenced using either SFB (grey) or LFB (green) size selection. For each sample, the same multiplex PCR products were split and sequenced twice, on two separate Flongle Flow Cells, using either SFB or LFB. Key difference is the *P. falciparum* mapping percentage, indicated at right.

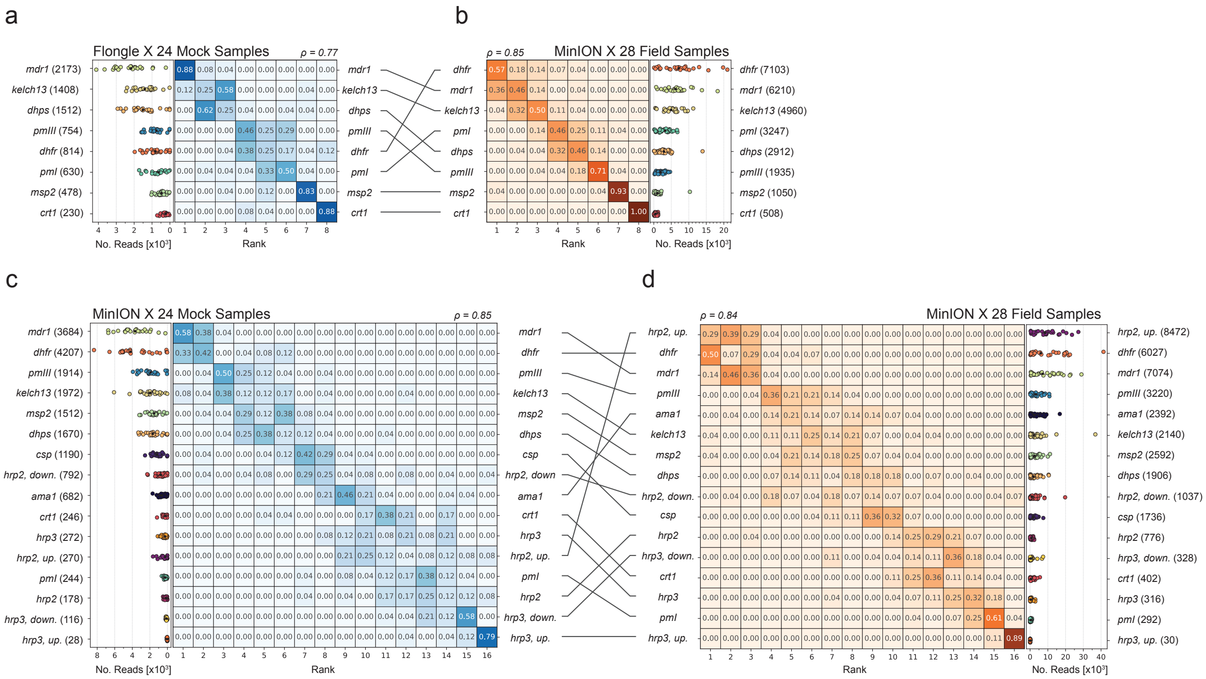


**Supplementary Figure 4: Order of amplicon abundances for the NOMADS8 and NOMADS16 panels in mock and field samples**. (a) NOMADS8 panel data for 24 mock samples sequenced on a Flongle Flow Cell. Left pane displays scatter plot of amplicon abundances (x-axis) for each target (y-axis). Median number of reads per amplicon indicated in brackets next to amplicon target name. Right pane displays heatmap with amplicons as rows and their rank-order by abundance as columns. Each cell indicates the fraction of samples in which a given target amplicon had a given abundance rank. Spearman's $\rho$ for amplicon abundances across samples shown above heatmap. Amplicons are ordered on y-axis by their median ranking. (b) Same as (a) but for 28 field samples sequenced on a MinION. Between (a) and (b), lines connect targets showing variation in abundance between mock and field samples. (d) and (c) same as (a) and (b), but for the NOMADS16 panel.

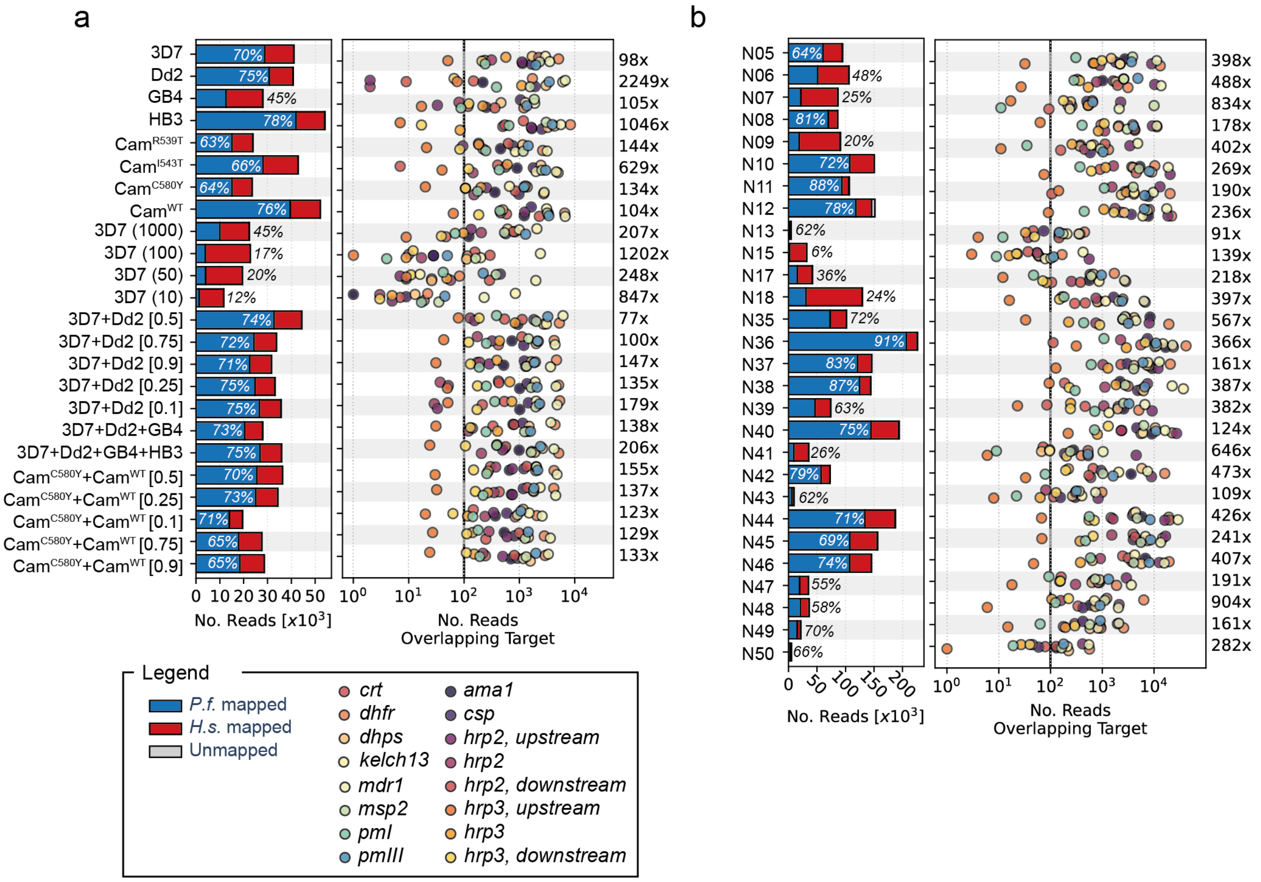


**Supplementary Figure 5: Sequencing throughput and coverage across samples and target genes for the NOMADS16 panel.** (a) Sequencing results for NOMADS16 on 24 mock samples sequenced on a standard MinION Flow Cell (R9.4.1). Bar plot (left pane) displays the total number of reads generated for each sample stratified by mapping outcome: mapped to *P. falciparum* (P.f.) (blue), human (H.s.) (red), or failing to map (grey, too few to be visible). *P.f.* mapping percentages indicated with text. Scatter plot (right pane) displays the number of reads overlapping each target gene (labeled by colors) after mapping for each sample. Note number of reads (x-axis) is displayed in log-scale. For most samples, majority of target genes have >100X coverage. Number at right (e.g. 98X for 3D7) gives the fold-difference between the highest coverage and lowest coverage target (see Methods). (c) Same as (b) but for 28 field samples collected as DBS from Kaoma, Zambia.

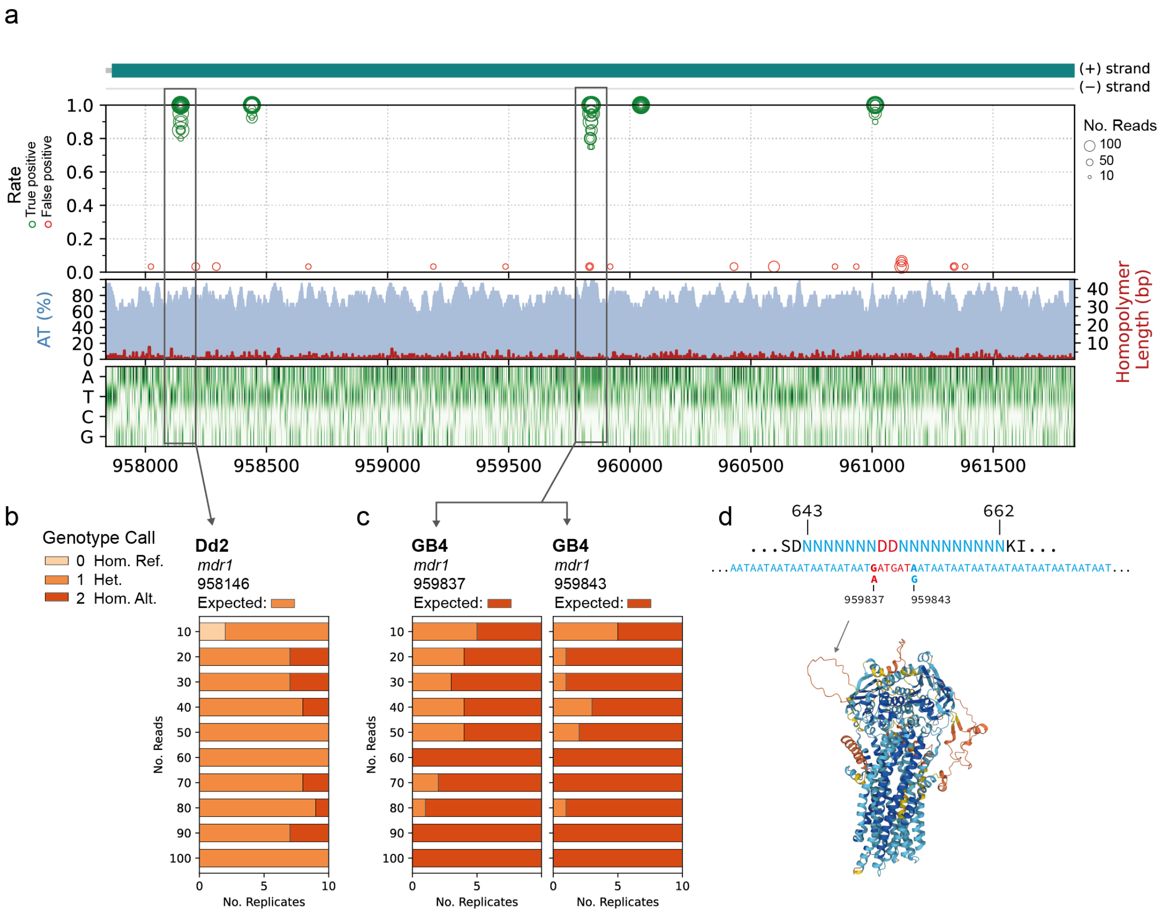

**Supplementary Figure 6: SNP calling errors in *mdr1* are have two primary sources.** (a) Diagram of *mdr1* amplicon region showing true positive and false positive rates for *in silico* replicates of Dd2, GB4 and HB3. From top to bottom, panels show an exon diagram of *mdr1*; the true positive rate (green) and false positive rate (red) of each site across thirty replicates at a given read depth (indicated by circle size); (A+T) % in 20bp sliding windows (blue shade) and homopolymer length (red line); and heatmap of nucleotide composition. Squares highlight high error regions at chromosomal position 958146 and positions 959837 and 959837. (b) Genotype calling results from `Clair3` for the 958146 site in the Dd2 mock sample. Barplots show genotype call (color) across the ten *in silico* replicates produced at each read depth. In some replicates, the site is erroneously called as homozygous alternative (Hom. alt), rather than heterozygous (Het.). (c) Same as (b) but for sites 959837 and 959837 in GB4. Here, there are erroneous heterozygous calls in some replicates. (d) Amino acid and nucleotide sequence surrounding sites 959837 and 959843, which fall within a repetitive region of 7xAsn, 2xAsp, 10xAsn. Below *mdr1* structure as predicted by AlphaFold. Codons 643 to 662 form an unstructured loop, indicated with an arrow.

| Step | Item | Supplier | Item Code | Cost per Item (USD) | Cost per Sample (USD) |
|---|---|---|---|---|---|
| DNA Extraction | QIAamp DNA Micro Kit (50) | Qiagen | 56304 | $316.00 | $6.32 |
| sWGA | phi29 DNA Polymerase | NEB | M0269L | $251.00 | $2.01 |
| sWGA | 1000uM sWGA Primer Mix | IDT | | | $- |
| sWGA | NEB 10mM dNTP Solution Mix | NEB | N0447L | $274.00 | $0.34 |
| Multiplex PCR | NOMADS16 Primers | IDT | | | $- |
| Multiplex PCR | KAPA HIFI + dNTPS (100U) | Roche | KK2101 | $83.00 | $0.83 |
| Barcoding | Qubit™ dsDNA HS Assay Kit (500) | ThermoFisher | Q32854 | $378.00 | $0.76 |
| Barcoding | Native Barcoding Expansion 96 | ONT | EXP-NBD196 | $1,240.00 | $0.16 |
| Barcoding | NEBNext UltraII DNA Library Prep with SPRI | NEB | E7103L | $2,514.00 | $5.24 |
| Adapter Ligation | Ligation Sequencing Kit | ONT | SQK-LSK109 | $599.00 | $1.04 |
| Adapter Ligation | NEBNext Quick Ligation Module | NEB | E6056L | $1,376.00 | $0.29 |
| Sequencing | R9.4.1 Flow Cell | ONT | FLO-MIN106D | $675.00 | $7.03 |
| | | | | **TOTAL** | **$24.01** |

**Supplementary Table 1: Consumable costs for amplicon-based nanopore sequencing on a standard MinION Flow Cell (FLO-MIN106D).** From barcoding step onwards, all per sample costs are calculated assuming 96 samples are run per R9.4.1. Flow Cell, and no washes are performed. The R9.4.1. Flow Cell cost is the individual cost of a Flow Cell purchased as a pack of 24. Primers contribute less than USD $0.01 to per sample cost and so are excluded. The remaining USD $1 per sample absorbs the cost of plasticware, nuclease-free water, ethanol, etc. Flongle Flow Cell (FLO-FLG001) decreases costs of sequencing, but increases cost of adapter ligation; as only 24 samples are prepared per ligation reaction. Costs were retrieved from supplier websites on 2023/01/30.

| No. | Sample Name | Strains | COI | Proportions | Parasitemia (p/uL) |
|---|---|---|---|---|---|
| 1 | 3D7 | 3D7 | 1 | 1 | 10,000 |
| 2 | Dd2 | Dd2 | 1 | 1 | 10,000 |
| 3 | GB4 | GB4 | 1 | 1 | 10,000 |
| 4 | HB3 | HB3 | 1 | 1 | 10,000 |
| 5 | $Cam^{R539T}$ | IPC 5202 | 1 | 1 | 10,000 |
| 6 | $Cam^{I543T}$ | IPC 4912 | 1 | 1 | 10,000 |
| 7 | $Cam^{C580Y}$ | IPC 3445 | 1 | 1 | 10,000 |
| 8 | $Cam^{WT}$ | IPC 3663 | 1 | 1 | 10,000 |
| 9 | 3D7 (1,000) | 3D7 | 1 | 1 | 1,000 |
| 10 | 3D7 (100) | 3D7 | 1 | 1 | 100 |
| 11 | 3D7 (50) | 3D7 | 1 | 1 | 50 |
| 12 | 3D7 (10) | 3D7 | 1 | 1 | 10 |
| 13 | 3D7+Dd2 [0.5] | 3D7; Dd2 | 2 | 0.5:0.5 | 10,000 |
| 14 | 3D7+Dd2 [0.75] | 3D7; Dd2 | 2 | 0.75:0.25 | 10,000 |
| 15 | 3D7+Dd2 [0.9] | 3D7; Dd2 | 2 | 0.9:0.1 | 10,000 |
| 16 | 3D7+Dd2 [0.25] | 3D7; Dd2 | 2 | 0.25:0.75 | 10,000 |
| 17 | 3D7+Dd2 [0.1] | 3D7; Dd2 | 2 | 0.1:0.9 | 10,000 |
| 18 | 3D7+Dd2+GB4 | 3D7; Dd2; GB4 | 3 | 0.33:0.33:0.33 | 10,000 |
| 19 | 3D7+Dd2+GB4+HB3 | 3D7; Dd2; GB4; HB3 | 4 | 0.25:0.25:0.25:0.25 | 10,000 |
| 20 | $Cam^{C580Y}$+$Cam^{WT}$ [0.5] | IPC 3445; IPC 3663 | 2 | 0.5:0.5 | 10,000 |
| 21 | $Cam^{C580Y}$+$Cam^{WT}$ [0.25] | IPC 3445; IPC 3663 | 2 | 0.25:0.75 | 10,000 |
| 22 | $Cam^{C580Y}$+$Cam^{WT}$ [0.1] | IPC 3445; IPC 3663 | 2 | 0.1:0.9 | 10,000 |
| 23 | $Cam^{C580Y}$+$Cam^{WT}$ [0.75] | IPC 3445; IPC 3663 | 2 | 0.75:0.25 | 10,000 |
| 24 | $Cam^{C580Y}$+$Cam^{WT}$ [0.9] | IPC 3445; IPC 3663 | 2 | 0.9:0.1 | 10,000 |

**Supplementary Table 2: Mock samples used throughout in this study**. Mock samples containing an *in vitro* mixture of genomic DNA from different *P. falciparum* laboratory/cultured strains and human DNA (Methods). For sample names, normal parentheses enclose parasitemia of the mock sample; when absent the parasitemia is 10,000p/$\mu$L. Similarly, square parentheses enclose the proportion of the first strain listed in the mock sample, for cases where COI> 1. For example '3D7+Dd2[0.25]' indicates that 3D7 is at a proportion of 0.25. When square brackets are empty, all samples are at balanced proportions.

| Sample | No. Reads | Chrom. | Pos. | Ref | Alt | True GT | Sample GT | n | N |
|---|---|---|---|---|---|---|---|---|---|
| GB4 | 50 | Pf3D7_05_v3 | 959837 | G | A | 2 | 1 | 4 | 10 |
| GB4 | 50 | Pf3D7_05_v3 | 959843 | A | G | 2 | 1 | 2 | 10 |
| HB3 | 50 | Pf3D7_05_v3 | 961013 | A | G | 2 | 1 | 1 | 10 |
| Dd2 | 60 | Pf3D7_05_v3 | 961122 | G | A | 0 | 1 | 1 | 10 |
| Dd2 | 70 | Pf3D7_05_v3 | 958146 | A | T | 1 | 2 | 2 | 10 |
| GB4 | 70 | Pf3D7_05_v3 | 959837 | G | A | 2 | 1 | 2 | 10 |
| Dd2 | 80 | Pf3D7_05_v3 | 958146 | A | T | 1 | 2 | 1 | 10 |
| GB4 | 80 | Pf3D7_05_v3 | 959837 | G | A | 2 | 1 | 1 | 10 |
| GB4 | 80 | Pf3D7_05_v3 | 959843 | A | G | 2 | 1 | 1 | 10 |
| Dd2 | 90 | Pf3D7_05_v3 | 958146 | A | T | 1 | 2 | 3 | 10 |

**Supplementary Table 3: All SNP errors observed in *mdr1* in mock samples that have been downsampled to fifty reads or greater**. For a given substitution error, the true genotype ('True GT') and observed genotype in the sample ('Sample GT') are shown, where 0 indicates homozygous reference; 1 indicates heterozygous; and 2 indicates homozygous alternative. The number of downsampled replicates with the error is given in column 'n'. The total number of downsampled replicates for that sample and number of reads in column 'N'.
